# Ovipositor and mouthparts in a fossil insect support a novel ecological role for early orthopterans in Pennsylvanian forests

**DOI:** 10.1101/2021.06.18.448941

**Authors:** Lu Chen, Jun-Jie Gu, Qiang Yang, Dong Ren, Alexander Blanke, Olivier Béthoux

**Affiliations:** College of Life Sciences and Academy for Multidisciplinary Studies, Capital Normal University, 105 Xisanhuanbeilu, Haidian District, Beijing 100048, China; Institute of Ecological Agriculture, College of Agronomy, Sichuan Agricultural University, Chengdu, Sichuan, 611130, China; School of Life Sciences, Guangzhou University, 230 Waihuanxi Road, Guangzhou Higher Education Mega Center, Guangzhou 510006, China; Institute of Evolutionary Biology and Animal Ecology, University of Bonn, An der Immenburg 1, 53121 Bonn, Germany; CR2P (Centre de Recherche en Paléontologie – Paris), MNHN – CNRS – Sorbonne Université; Muséum National d’Histoire Naturelle, 57 rue Cuvier, CP 38, 75005 Paris, France

## Abstract

Lobeattid insects represented a high portion of the earliest known, Pennsylvanian insect faunas. However, their systematic affinities and their role as foliage feeders which severely influenced their ecosystems remain debated. We investigated hundreds of samples of a new lobeattid species from the Xiaheyan locality using Reflectance Transforming Imaging combined with geometric morphometrics in order to assess its morphology, infer its ecological role, and phylogenetic position. *Ctenoptilus frequens* sp. nov. possessed a sword-shaped ovipositor whose valves interlocked by two ball-and-socket mechanisms. This unambiguously supports lobeattids as stem-relatives of all Orthoptera (crickets, grasshoppers, katydids). Given the herein presented and other remains, it follows that this group experienced an early diversification coupled with high levels of abundance. The ovipositor shape additionally indicates that ground was the preferred substrate for eggs. Visible mouthparts made it possible to assess the efficiency of the mandibular food uptake system in comparison to a wide array of recent species. The new species was omnivorous which explains the paucity of external damage on contemporaneous plant foliage.

## INTRODUCTION

The earliest known insect faunas in the Pennsylvanian, ca. 307 million years ago, were populated by species displaying mixtures of inherited (plesiomorphic) and derived (apomorphic) conditions, such as the griffenflies, but also by highly specialized groups, such as the gracile and sap-feeding megasecopterans. A prominent portion of this past insect fauna were the so-called ‘lobeattid insects’. They have been recovered from all major Pennsylvanian outcrops, where some species can abound (*1-3*). Indeed, at the Xiaheyan locality, China, for which quantitative data are available, they collectively account for more than half of all insect occurrences (*4*). Additionally, another extinct group - the order Cnemidolestodea - composed of derived relatives of lobeattid insects, was likewise ubiquitously distributed during the Pennsylvanian until the onset of the Permian (*5*).

The systematic affinities of this abundant fraction of the earliest insect faunas are debated. They have been regarded as stem-relatives of either Orthoptera (*6*) or of other lineages within Polyneoptera (*7, 8*). A core point of the debate is the presumed wing venation groundplan of insects, which, however, will remain elusive until Mississipian or even earlier fossil wings are discovered. Ecological preferences of lobeattid insects are also poorly known. They have been traditionally regarded as foliage feeders (*9*) but, given their abundance, this stands in contrast to the paucity of documented foliage external damage during that time.

The Xiaheyan locality is unique in several respects (*4*), including the amount of insect material it contains. Over the past decade a collection of several thousand specimens was unearthed, allowing for highly detailed analyses of e.g. ovipositor and mouthparts morphology of extinct insect orders (*10*). These character systems are investigated herein in a new lobeattid species, documented based on an abundant material using Reflectance Transforming Imaging (RTI). Dietary preferences were inferred based on a comparative geometric morphometric analysis relying on an extensive dataset of extant polyneopteran species, with a focus on Orthoptera. Together, this investigation provides decisive information regarding the dietary niche of lobeattid insects and their phylogenetic affinities.

## RESULTS

### Systematic Palaeontology

Archaeorthoptera Béthoux and Nel, 2002

Ctenoptilidae Aristov, 2014

*Ctenoptilus* Lameere, 1917

*Ctenoptilus frequens* Chen, Gu & Béthoux, sp. nov.

**LSID (Life Science Identifier)**. xxxx

**Etymology**. Based on ‘*frequens*’ (‘frequent’ in Latin), referring to the abundance of the species at the Xiaheyan locality.

**Holotype**. Specimen CNU-NX1-326 (female individual; Fig. 1).

**Figure 1.**
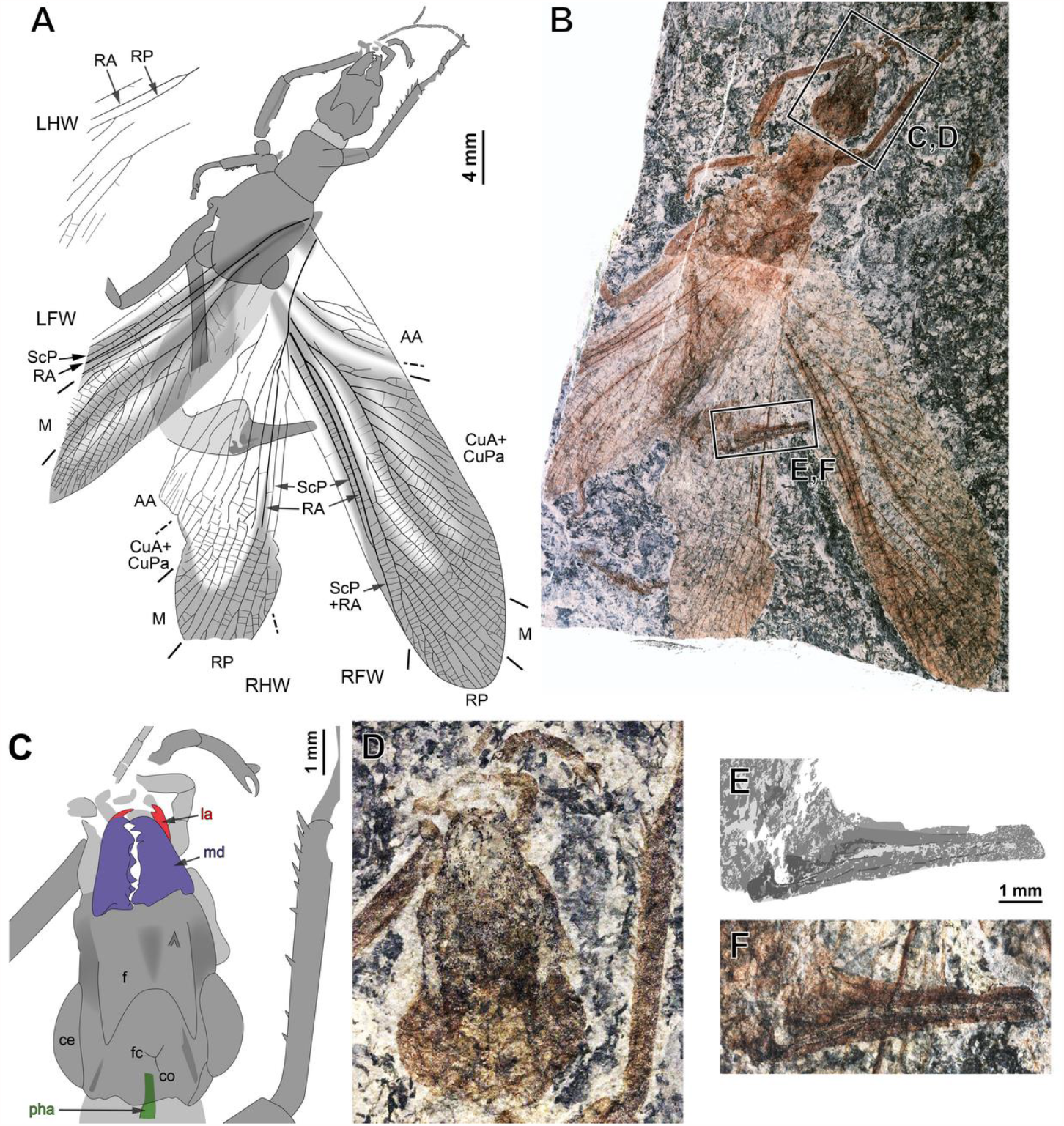
Ctenoptilus frequens sp. nov., holotype (CNU-NX1-326). (A) habitus drawing and (B) habitus photograph (composite); (C–D) details of head and right foreleg (location as indicated in B), (C) color-coded interpretative drawing and (D) photograph (composite); and (E–F) details of ovipositor (location as indicated in B), (E) drawing and (F) photograph (composite). Color-coding and associated abbreviations: red, lacina (la); dark blue-purple, mandible (md); green, pharynx (pha). Other indications, head: ce, composite eye; f, frons; co, coronal cleavage line; fc, frontal cleavage line. Wing morphology abbreviations: LFW, left forewing; LHW, left hind wing; RFW, right forewing; RHW, right hind wing; ScP, posterior Subcosta; RA, anterior Radius; RP, posterior Radius; M, Media; CuA, anterior Cubitus; CuPa, anterior branch of posterior Cubitus; CuPb, posterior branch of posterior Cubitus; AA, anterior Analis.

**Referred material**. See Supplemental Text section 2.1.2.

**Locality and horizon**. Xiaheyan Village, Zhongwei City, Yanghugou Formation (Ningxia Hui Autonomous Region, China); latest Bashkirian (latest Duckmantian) to middle Moscovian (Bolsovian), early Pennsylvanian (*4*).

**Differential diagnosis**. The species is largely similar to *Ctenoptilus elongatus* (Brongniart, 1893), in particular in its wing venation (Supplemental Text section 2.1.2). However, it differs from it in its smaller size (deduced from forewing length) and its prothorax longer than wide (as opposed to quadrangular).

**General description**. See Supplemental Text section 2.1.2.

**Specimens description**. See Supplemental Text section 2.1 and figs. S2–S8; details of ovipositor, see Fig. 2; details of head, see Fig. 3.

**Figure 2.**
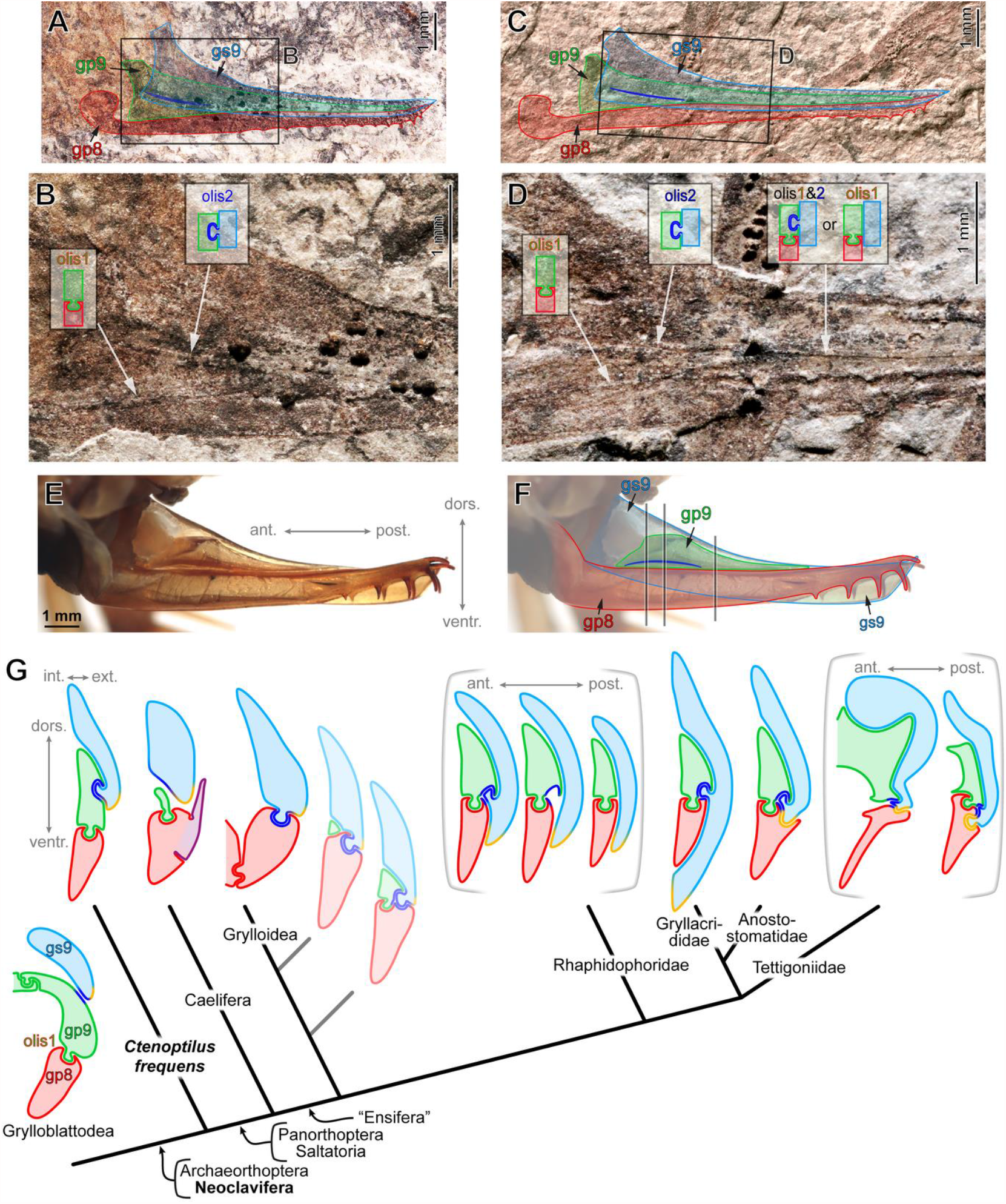
External ovipositor in laterial view, (A–D) Ctenoptilus frequens sp. nov. and (E–F) Ceuthophilus sp. (Orthoptera: Rhaphidophoridae; extant), and (G) ovipositor cross-sections in Grylloblattodea, *Ctenoptilus frequens* sp. nov. and a selection of extant Orthoptera possessing well-developed ovipositors (schemes, left side, not to scale; and see Supplemental Text section 2.2), mapped onto a consensual phylogeny. (A–B) specimen CNU-NX1-749, (A) overview with interpretation (and see Figure S7A–C) and (B) detail of base as located on A; (C–D) specimen CNU-NX1-742, (C) overview with interpretation (and see Figure S8) and (D) detail of base as located on C; (E) General view (left side, flipped horizontally, left gs9 removed) and (F) the same, annotated (the three black vertical lines indicate the position of the three schematic sections represented in G, Rhaphidophoridae); (G) pale cross-sections along the stem of Grylloidea hypothetical; sections delineated by brackets represent conditions along the antero-posterior axis of a single type. Color-coding and associated abbreviations: light blue, gonostylus IX (gs9); light green, gonapophysis IX (gp9); red, gonapophysis VIII (gp8); royal blue, secondary olistheter (olis2); light orange, tertiary olistheter (olis3); purple, ‘lateral basivalvular sclerite’ (specific to Caelifera). Other indications: olis1, primary olistheter; int./ext., internal/external, respectively; dors./ventr., dorsal/ventral, respectively; and ant./post., anterior/posterior, respectively.

**Figure 3.**
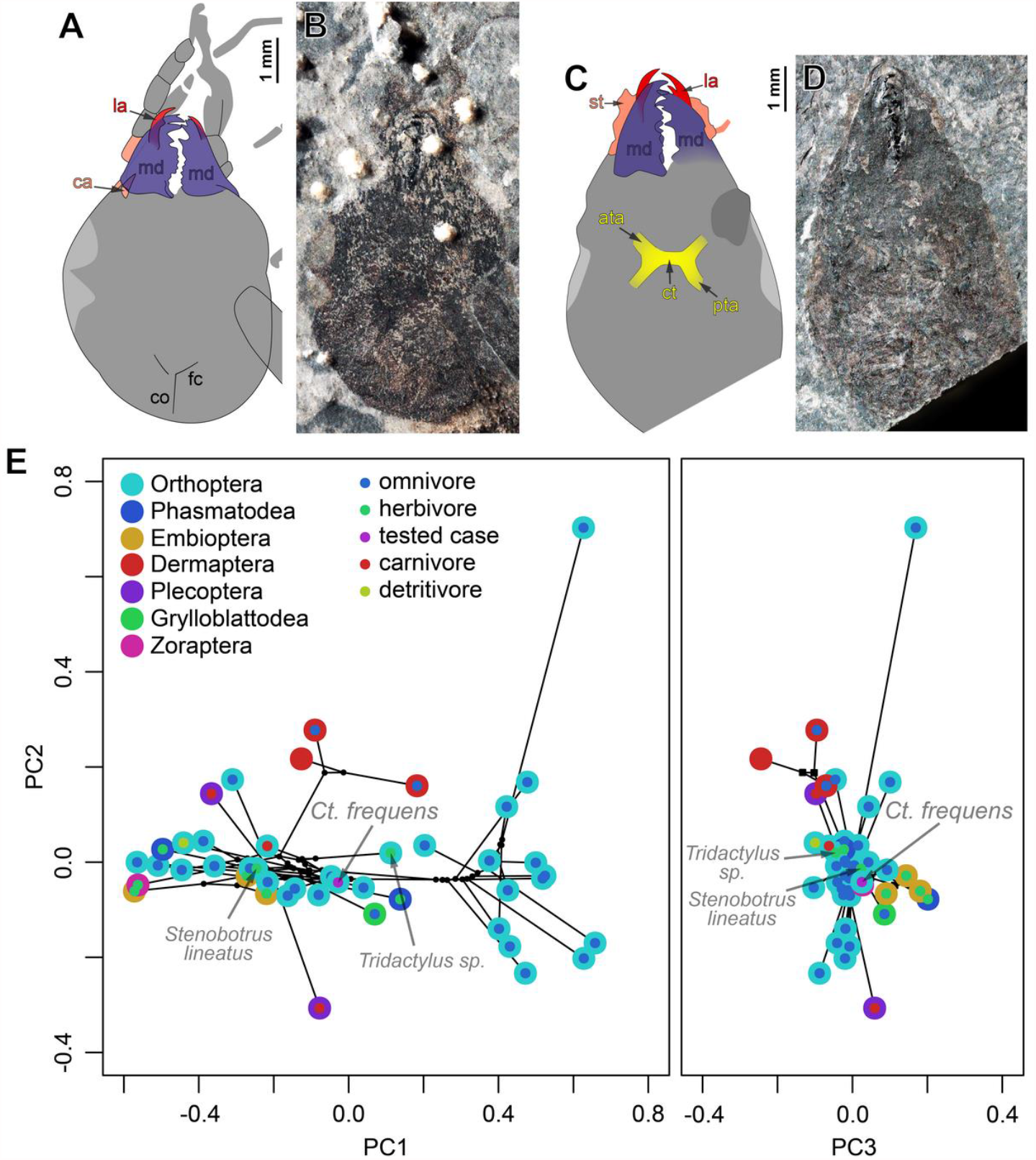
Head morphology (A–D) in Ctenoptilus frequens sp. nov. and (E) mandibular mechanical advantage in Ctenoptilus frequens sp. nov. and a selection of polyneopteran species. (A–B) Specimen CNU-NX1-754, (D) color-coded interpretative drawing and (E) photograph (composite) (as located on Figure S7I); (C–D) Specimen CNU-NX1-764, (D) color-coded interpretative drawing and (E) photograph (composite); (E) Principal component analysis of the mandibular mechanical advantage. Color-coding: (A– D) red, lacina (la); salmon, cardinal and stipital sclerites (ca and st, respectively); dark blue-purple, mandible (md); yellow, tentorium, including anterior tentorial arm (ata), posterior tentorial arm (pta) and corpotentorium (ct). Other indications: co, coronal cleavage line; fc, frontal cleavage line.

### Ovipositor morphology

The external genitalia in insects consist primarily of a pair of mesal extensions, the so-called *gonapophyses*, or ovipositor blades, and a pair of lateral projections, the so-called gonostyli, or ovipositor sheaths on abdominal segments 8 and 9. These sclerotized elements are collectively referred to as ‘valves’. The studied fossils possess three pairs of valves in their ovipositor, each strongly sclerotized (Fig. 2A–D, and figs. S7B and C and S8B and C). Especially the valve margins are still visible in the anterior area (‘base’), including the dorsal margin of the gonostylus 9 (gs9), the ventral margin of the gonapophysis 9 (gp9), and the dorsal and ventral margins of gonapophysis 8 (gp8). All observed ovipositors, but in particular the one of specimen CNU-NX1-742 (Fig. 2C and D, and fig. S8B and C), display, from the second third of their length onwards, a thin longitudinal line much sharper and more developed than other visible linear structures in the area. This is the primary olistheter (olis1), a tongue like structure which commonly interlocks gp9 and gp8 in extant insects having more or less well-developed external ovipositors (Fig. 2G) (*11*). In the distal half of the ovipositor, the linear structure occurring between the dorsal edge of gs9 and olis1 is interpreted as the dorsal margin of gp9.

Together with the position of the antero-basal apophysis (=outgrowth) of this valve, the anterior margin of gp9 can then be traced. The extent of olis1 indicates that gp9 reaches the ovipositor apex, which is corroborated by the length of its inferred dorsal margin, well visible in specimen CNU-NX1-749 (Fig. 2A and B, and fig. S7B and C). This specimen also shows that gp8 bears ventrally-oriented teeth, more prominent and densely distributed near the apex similar to recent orthopterans (Fig. 2). The location of the dorsal margin of gp9 could not be observed with confidence near the base, which might be due to a lower degree of sclerotization.

This morphology implies that, at the base, dorsal to the anterior margin of gp8, only gs9 and gp9 occur. Therefore, the sharp and heavily sclerotized longitudinal line, located slightly dorsally with respect to the ventral margin of gs9, can only be an olistheter interlocking these two valves (olis2). This second olistheter reaches olis1 but its development beyond this point could not be inferred with the available material. The occurrence of a mechanism locking gs9 onto gp9 is further supported by the fact that these valves remained connected to each other in the specimen CNU-NX1-742 even though it endured heavy decay (head and ovipositor detached from thorax and abdomen, respectively; fig. S8). The ovipositor configuration in *Ct. elongatus* conforms the one assumed for the Orthoptera groundplan, as exemplified by extant Raphidophoridae (Fig. 2E and F, and see Fig. G; and Supplemental Text section 2.2) in which a second olistheter occurs in addition to olis1 and interlocks gs9 and gp9.

### Mandibular mechanical advantage

The head and mouthpart morphology could be investigated in more detail in six specimens (see supplementary text) while we could study the mechanical advantage (MA; see section 1.4 of the supplementary text) of their mandibles in four of the six (viz. CNU-NX1-326, −747, −754, −764). The MA is defined as the inlever to outlever ratio and thus indicates the percentage of force transmitted to the food item (i.e. the effectivity of the lever system). Therefore, the MA allows for a size independent comparison of the relative efficiencies of force transmission to the food item. Low MA values usually indicate quick biting with low force transmission typical for predators, while high MA values indicate comparatively slow biting with higher force transmission typical for non-predatory species.

Calculation of the mechanical advantage along the entire gnathal edge revealed characteristic MA curve progressions for the studied taxa (Supplemental Text section 2.3, and fig. S9). Compared to the studied fossils, recent Dermaptera, Embioptera, and Phasmatodea showed comparatively high MAs with an almost linear curve progression towards more distal parts of the mandibular incisivi whereas Plecoptera, Zoraptera, and Grylloblattodea were located at the lower end of the MA range with a gently exponential decrease towards the distal incisivi. The analysed extant Orthoptera occupy a comparatively wide functional space, with lineages at the higher and lower end of the MA range. The composite fossil mandible representation (CFMR) of *Ct. frequens* (see Methods) is located in the centre of the observed range of MAs for Orthoptera (Fig. 3).

A polynomial function of the fifth order resulted in the best relative fit on the MA curves according to the AIC value (−661.3, see Methods). The five common coefficients were subjected to a principal component analysis (PCA, Fig. 3E), and, because phylogenetic signal was detected (K = 1.03316; p = 0.0001), also analysed using a phylogenetic principal component analysis (pPCA) (Supplemental Text section 2.3, and fig. S10). The first four principal components (PCs) accounted for 96.8% (PCA) / 96% (pPCA) of the variation in MA (tab. S3).

In both PCAs, PC1 mainly codes for the vertical position of the MA curve, i.e. the effectivity of the force transmission along the whole toothrow, while PC2 mainly codes for the curvature, i.e. whether there is an almost linear or a gently exponential decrease in the effectivity of force transmission. Due to the narrow distribution of species along PC3 it was not possible to associate a clear biomechanical pattern to this PC.

The CFMR of *Ct. frequens* is located at the centre of the first three PCs (Fig. 3E). Omnivorous Orthoptera and all herbivore taxa, with the exception of *Apotrechus*, are located along the width of PC1, while there is a tendency for the carnivorous taxa within the sampling to be spread along PC2.

## DISCUSSION

### Ovipositor morphology and phylogeny

Our analysis of material of *Ct. frequens* provides unequivocal evidence that an olis2 occurs in this species. Therefore, the new species was an orthopteran. Owing to the lack of jump-related specialisation in the hind leg, the species can be excluded from Saltatoria. Due to its wing morphology, with a CuPa vein lacking a fork basal to its fusion with the CuA vein, the new species can also be excluded from the Panorthoptera. It follows that *Ct. frequens*, and its various Pennsylvanian relatives collectively referred to as ‘lobeattid insects’, are stem-relatives of Orthoptera (Fig. 2G).

The organizational diversity of elements which form the external ovipositor in Orthoptera made it difficult to reconstruct its evolution based on extant species only (*12-16*; Supplemental Text section 2.2). Comparison has traditionally been made between Grylloblattodea (rock-crawlers) and Orthoptera (*15*) even though the two groups are not closely related. In both groups the ovipositor displays an elongate gonostylus IX (gs9) and a ball-and-socket locking mechanism, the so-called primary olistheter (olis1), interlocking gonapophyses IX onto gonapophyses VIII (gp9 and gp8, respectively; Fig. 2G). This primary olistheter occurs widely among insects (*11*). Orthoptera possess a variety of additional olistheters, including one interlocking gs9 onto gp9 (royal blue in Fig. 2; olis2), as exemplified by Rhaphidophoridae (cave crickets; Fig. 2E and F) and Gryllacrididae (raspy & king crickets). The occurrence of an olis2 is diagnostic of ensiferan (‘sword-bearing’) Orthoptera (*14*; and see below).

Even though it is unclear how far posteriorly olis2 extends in *Ct. frequens*, the asserted phylogenetic placement of this species provides new insights on the evolution of ovipositor interlocking mechanisms in Orthoptera (*14*; Fig. 2G). The ovipositor interlocking mechanisms is comparable to the one of Rhaphidophoridae, the main differences concern the rachis (‘ball’ as in ‘ball-and-socket’), which is limited to a short protrusion in these insects, while the aulax (‘socket’ as in ‘ball-and-socket’) extends further posteriorly. In addition, gs9 extends more ventrally, concealing gp8 for some distance. Compared to Gryllacrididae the only notable difference in *Ct. frequens* is the ventral extension of gs9 in the former. In Anostostomatidae the ventral margin of gs9 enters a socket in gp8, regarded as composing the premises of a third olistheter (olis3). This new structure ultimately replaces olis2 in Tettigoniidae and allows a coupling of gs9 with gp8.

Grylloidea (true crickets) and *Ct. frequens* are separated by more severe morphological differences. In the former, gp9 is reduced or even absent, and an olistheter connects gs9 and gp8.

Unlike other orthopterans displaying a well-developed external ovipositor, Caelifera (grasshoppers) use valves for digging a tunnel to accommodate their entire abdomen and, additionally, dig egg pods (*17-19*). The shoving operation to move forward is accomplished by powerful, rhythmic, dorso-ventral openings and closings of two sets of valves (*16*), gs9 and gp8+gp9, the two latter ones being interlocked via olis1. Even though gp9 is often reduced, it plays an important role in the closing of the ovipositor via muscles attached to it (*16*). Obviously, an olistheter interlocking gs9 and gp8 (i.e. olis2) would impede such movements. Given the ovipositor configuration and phylogenetic placement of *Ct. frequens*, it follows that the olis2 was lost in Caelifera, a likely consequence of their highly derived oviposition technique.

The evolutionary scenario resulting from our findings in *Ct. frequens* addresses a long-standing debate on the respective position of the two sub-orders of Orthoptera, Ensifera and Caelifera. On the basis of early, fossil Saltatoria displaying elongate ovipositors, palaeontologists already assumed that caeliferans derived from ensiferans (*20*). However, the placement of these fossils remained contentious, leaving it possible that both sub-orders derived from an earlier, unspecialized assemblage (*21*). The discovery of an elongate ovipositor in the stem-orthopteran *Ct. frequens* provides a definitive demonstration that caeliferans derived from ensiferans. Because rock-crawlers can also be understood as possessing an elongate ovipositor, which would render the term ‘Ensifera’ ambiguous, it is proposed to coin a new taxon name, Neoclavifera, to encompass species bearing an olis2, i.e. all extant orthopterans and their known stem-relatives (Fig. 2G; Supplemental Text section 2.1.1).

Another important input on the early evolution of orthopterans regards the abundance of lobeattids. Indeed, these insects are emerging as the main component of Pennsylvanian insect faunas. They have been reported in high numbers from all major Pennsylvanian deposits (*1-3*; and Supplemental Text section 2.1), such as *Miamia bronsoni* at Mazon Creek (*3*). At Xiaheyan, they collectively account for more than half of all insect occurrences (*4*). Besides a high abundance, lobeattids and other stem-orthopterans compose a species-rich group at Xiaheyan, where they represent about a third of all insect species currently known to occur at this locality (Supplemental Text section 3, taxon Archaeorthoptera). Orthoptera, which represent the bulk of extant polyneopteran insect diversity, therefore must have diversified early during their evolution.

### Ovipositor shape and use

Extant Orthoptera resort to a wide diversity of substrates where to lay eggs, including ground, decaying leaves or wood, and stems or leaves of living plants (*12, 13, 22, 23*). This operation aims at ensuring a degree of moisture conditions suitable for eggs to fully develop, and providing protection, for example against predation. Ground is the preferred substrate of the majority of Orthoptera, including Caelifera (*18, 19, 24*; and see above). Within this group, the epiphytic and endophytic habits of several, inner lineages represent derived conditions (*25, 26*). This habit translates into finely serrated ovipositor valves, including gs9.

As for ‘ensiferan’ Orthoptera, they generally possess a pointed and elongate ovipositor used to insert eggs in various substrates. In Grylloidea females insert eggs in the ground using a needle-like ovipositor, or deposit them in subterranean chambers or burrows adults may inhabit, in which case the ovipositor is usually reduced (*13, 27, 28*). However, within Grylloidea, three groups, the Trigonidiinae, the Aphonoidinii and the Oecanthinae, evolved oviposition in plants. In the former, which lay eggs in soft plant material, gs9 displays serration in its distal third, along its dorsal edge (*29, 30*). In contrast, both Aphonoidinii and Oecanthinae lay eggs in more robust plant material, translating into apices of gs9 provided with strongly sclerotized sets of teeth and hooks (*27*). In Oecanthinae, in which oviposition functioning was studied in most detail, the alternate back and forth movements of gp8 induce apices of gs9 to alternately approximate and diverge (*31*), and therefore act as a shoving tool.

The Rhaphidophoridae commonly lay eggs into the ground, or, alternatively, into rotten leaves or wood (*32*). In the latter case the ovipositor is often curved. Interestingly, *Ceuthophilus* spp. use the ovipositor tip, somewhat truncated, to rake ground surface above oviposition holes (*33*), presumably to hide them. Anostostomatidae lay eggs in the ground or on walls of subterranean chambers (*34, 35*). These preferences also apply to both Gryllacrididae (*36, 37*) and Stenopelmatidae (*38*; not represented in Figure 2*G*), in which the ovipositor, if well-developed, is long, narrow, and rectilinear to curved (*39, 40*).

Although most Tettigoniidae (katydids) lay eggs in the ground, a variety of plant tissues, including galls, are also targeted by members of this very diverse family (*12, 41, 42*). As above, shape and serration relate, to a large extent, to the preferred substrate. A needle-shaped ovipositor generally indicates preference for ground, a sickle-shaped one for plant tissues. Curved ovipositors indicate preference for decaying wood, and more strongly falcate ones, which are usually also laterally flattened (as opposed to sub-cylindrical), preference for either bark crevices or leaf tissues. Katydids laying eggs in hollow grass stems or leaf sheaths possess straight to slightly falcate, flattened and unarmed ovipositors. Marked serration on the dorsal side of the ovipositor indicates preference for plant tissues.

Given the relation of ovipositor shape and substrate in recent species, *Ct. frequens*, with its needle-shaped ovipositor including ventrally oriented teeth, likely oviposited in the ground (Fig. 4). It is therefore unlikely that Pennsylvanian stem-orthopterans were responsible for endophytic oviposition traces documented for this epoch (*43, 44*).

**Figure 4.**
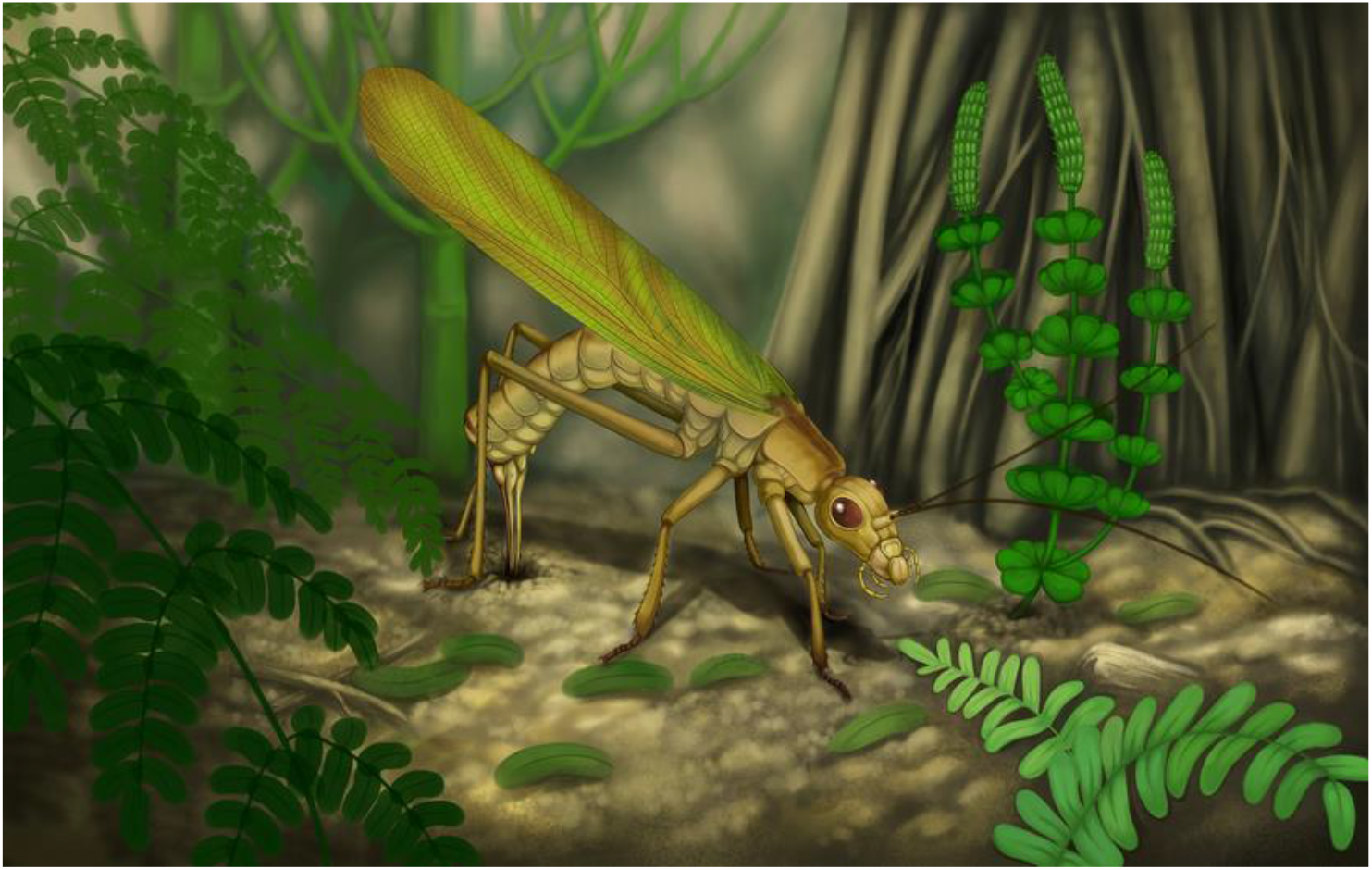
Reconstruction of a female of Ctenoptilus frequens sp. nov. laying eggs.

### Dietary preferences

Unlike in an extant tropical forest, a limited proportion of Pennsylvanian plant foliage experienced external damage, in particular generalized feeding types such as margin and hole feeding. Although such damages were reported from multiple localities, they are so rare that their occurrence was considered worth being reported (*44-47*). Quantitative data from Pennsylvanian localities indicate that generalized external damages were indeed rare, and concentrated on pteridosperms (‘seed ferns’; *48, 49*). Such damages have been traditionally assigned to Orthoptera and their purported stem-relatives (*9*). Indeed, investigation of mouthparts morphology in a subset of these insects suggested that, at least for the representatives belonging to the Panorthoptera/Saltatoria (Fig. 2G), these insects were herbivores (*50*). However, there is an inconsistency between the paucity of damage on Pennsylvanian plant foliage on the one hand, and the abundance of lobeattid insects on the other. If these insects were all external foliage feeders, evidence of such damage would be more prevalent.

Given the reconstruction of the mandibular gnathal edge and its position in PC space in relation to other Orthoptera and Polyneoptera (Fig. 3E; Supplemental Text section 2.3), *Ct. frequens* was likely an omnivore species - not a solely herbivorous or carnivorous one. The new species is the second most common insect species at Xiaheyan, where it occurs in all fossiliferous layers at a rate of ca. 10%. This implies that a significant portion of Pennsylvanian neopteran insects were opportunistic, omnivorous species, which reconciles the paucity of foliage damage with the abundance of stem-Orthoptera.

## MATERIAL AND METHODS

### Fossil material

The studied specimens are housed at the Key Laboratory of Insect Evolution and Environmental Changes, College of Life Sciences, Capital Normal University, Beijing, China (CNU). All specimens were collected from the locality near Xiaheyan village, where insect carcasses deposited in an interdeltaic bay (*4*).

The adopted morphological terminology is detailed in *SI Appendix* (section 1.1). Documentation methodology is detailed in *SI Appendix* (section 1.2.1). General habitus was investigated based on a selection of 23 specimens (including the holotype; Supplemental Text section 2.1.2). Ovipositor morphology was investigated based on four specimens (Supplemental Text section 1.2.2). Head and mouthparts morphology was investigated based on six specimens (Supplemental Text section 1.2.3).

### Comparative analyses

Fossil ovipositor morphology was compared to original material of extant species and to literature data (Supplemental Text sections 1.3.1, 2.2). Multiple interpretations of the fossil ovipositor morphology were considered. Among these, the favoured interpretation is the only one consistent with observations made on all specimens.

The mechanical advantage (MA) of the mandibles, i.e. the inlever to outlever ratio, indicates the effectivity of force transmission from the muscles to the food item (fig. S1). Apart from force transmission, the MA can also indicate the dietary niche and feeding habits (*51-53*). The MA was extracted from 43 recent polyneopteran species including 31 orthopterans and one composite fossil mandible representation (CFMR) of the newly described fossil species (Supplemental Text sections 1.3.2, 1.4, tab. S2). The CFMR was derived from a Procrustes superimposition (R package “geomorph” v.3.0.5; *54*) of four fossil specimens which showed low levels of overall distortion and a mandible orientation suitable for extraction of individual MAs. For comparison of species and inference of the dietary niche, a principal component analysis (PCA) and, due to the detection of significant phylogenetic signal, a phylogenetic PCA (pPCA; R package “phytools” v.0.6–44; *55*), were performed (for results of the pPCA see *SI Appendix* fig. S9, tab. S3).

## SUPPLEMENTAL INFORMATION

**Supplemental Information** includes Supplemental Text, ten Figures and three Tables and can be found at http://dx.doi.org/xxxx

**Supplemental data** (RTI files) are available for this paper at https://datadryad.org/stash/share/dmV-cfJHy2D475lLETIdQOzZ6HpxDWlnRk6xsw2yxXc

## Acknowledgements

We are grateful to the numerous students who collected fossil insects at Xiaheyan; to B. Kondratieff, S. Schoville and J. Lapeyrie for providing material for our comparative analysis of ovipositor morphology, and to V. Rommevaux for mounting and preparing this material; to S. Storozhenko for providing documentation; to S. Randolf for photographs of NHM Wien specimens; to S. Ingrish, C. Hemp and D. Rentz for discussion on ovipositor morphology in relation to substrate in ‘ensiferans’; to C. Labandeira for discussion on the intensity of folivory during the Pennsylvanian; and to D. Marjanovic and M. Laurin for discussion on nomenclatural procedures.

## Author contributions

D.R. and O.B. designed research; O.B. conducted fossil specimens preparation; L.C., A.B. and OB performed research and prepared figures; L.C., J.-J.G., Q.Y. and O.B. contributed fossil collects; A.B. and O.B. documented extant species; L.C., A.B., D.R. and O.B. wrote the paper.

## Data and material availability

All data needed to evaluate the conclusions in the paper are presented in the paper and/or the Supplemental Information, including Supplemental Text, ten figures and three tables (available at http://dx.doi.org/xxxx), and Supplemental data (available at https://datadryad.org/stash/share/dmV-cfJHy2D475lLETIdQOzZ6HpxDWlnRk6xsw2yxXc)

## Supplemental Information

### SUPPLEMENTAL TEXT

#### 1 Material and Methods

##### 1.1 Morphological terminology and abbreviations

###### 1.1.1 Head

**Ant**, antenna; **ata**, anterior tentorial arm; **ca**, cardinal sclerite; **ce**, compound eyes;**co**, coronal cleavage line; **ct**, corpotentorium; **f**, frons; **fc**, frontal cleavage line; **ga**, galea; **la**, lacinia; **il**, incisor lobe; **mo**, molar lobe; **md**, mandible; **mp**, maxillary palpus; **pha**, pharynx; **pta**, posterior tentorial arm; **st**, stipital sclerite.

###### 1.1.2 Wings

We use wing venation homologies proposed by Béthoux & Nel (*6*) for Archaeorthoptera. Corresponding abbreviations are: ScP, posterior Subcosta; R, Radius; RA, anterior Radius; RP, posterior Radius; M, Media; CuA, anterior Cubitus; CuP, posterior Cubitus; CuPa, anterior branch of CuP; CuPb, posterior branch of CuP; AA, anterior Analis; AA1, first anterior Analis; AA2, second anterior Analis. On figures, RFW, LFW, RHW and LHW refer to the left forewing, right forewing, right hind wing and left hind wing, respectively. A ‘furrow’ is a line along which veins and wing membrane are desclerotized. Median and cubital furrows commonly occur in insects.

###### 1.1.3 Ovipositor

Several terminologies have been used to refer to the elongate and sclerotized elements (herein collectively referred to as ‘valves’) composing the ovipositor in insects in general (tab. S1). We favoured Smith’s (*56*) terminology because it applies widely and is consensually admitted regarding homology hypotheses. In order to ease comparison we resorted to color-coding for selected structures, as follows: light blue, gonostylus (**gs9**); light green, gonapophysis IX (**gp9**); red, gonapophysis VIII (**gp8**); royal blue, olistheter 2 (**olis2**); light orange, olistheter 3 (**olis3**); purple, Ander’s (*57*) ‘lateral basivalvular sclerite’ (specific to Caelifera). Additional abbreviations applying to olistheter elements are as follows: olistheter 1 (**olis1**); **al**, aulax (i.e. groove, socket in ‘ball-and-socket’); **rh**, rhachis (i.e. ridge, ball in ‘ball-and-socket’).

##### 1.2 Documentation of fossil material

###### 1.2.1 General aspects

Handmade draft drawings were produced using a LEICA MZ12.5 dissecting microscope and illustrated with the aid of a drawing tube (Leica, Wetzlar, Germany). Photographs were taken using Canon EOS 550D or 5D Mark III digital cameras (Canon, Tokyo, Japan), coupled to a Canon 50 mm macro lens, a 100 mm macro lens, or a Canon MP-E 65 mm macro lens, all equipped with polarizing filters. Each specimen was photographed under dry condition and covered with a thin film of ethanol. When available, both imprints were photographed. These photographs were optimized using Adobe Photoshop CC 2015.5 (Adobe Systems, San Jose, CA, USA) and assembled, together with handmade drawings, into a single, multi-layered document. Reproduced photographs referred to as ‘composites’ are a combination of photographs of a dry specimen and the same under ethanol. In addition to traditional photographs, we computed Reflectance Transforming Imaging (RTI) files for details of several specimens. This corpus of data was used to produce illustrations using Adobe Illustrator CS6 (Adobe Systems, San Jose, CA, USA). Multi-layered documents (photographs only) and RTI files are provided in the associated Dryad dataset (*58*). Investigated specimens are listed in the section 2.1.2.

Measurements were based on complete specimens illustrated herein and are provided in the following format: minimum/average/maximum.

###### 1.2.2 Ovipositor morphology

The ovipositor morphology was investigated based on specimens CNU-NX1-326 [Fig. 1E and F; and see related files in Dryad repository (*58*)], −749 [Fig. 2A and B, and fig. S7B and C; and see related files in Dryad repository (*58*)], −742 [Fig. 2C and D, and fig. S8B and C; and see related files in Dryad repository (*58*)] and −743 [fig. S5O–Q; and see related files in Dryad repository (*58*)].

###### 1.2.3 Head and mouthparts morphology

The head and mouthpart morphology was investigated based on six specimens. Four of them (viz. CNU-NX1-326, −747, −754, −764) were investigated for the mechanical advantage (MA; see section 1.4) of their mandibles. The specimens CNU-NX1-749, and −756 were excluded from the analysis because their mandibles were preserved with a slight rotation in the frontal plane, this impeding an accurate measurement of the MA (see below).

##### 1.3 Documentation of extant material

###### 1.3.1 Ovipositor morphology

We complemented the available literature on the morphology of female terminalia which form the ovipositor in polyneopteran lineages and in Orthoptera in particular [*12-14, 57*; and see Klass (*11*) and references therein] by preparation of material belonging to various extant species (see section 2.2). External habitus was photographed under various angles. Terminalia, together with the ultimate abdominal segments, were then cut off and mounted in a polyester resin. Three to four sections were made at various levels and hand-polished. Direct observation and photographs (same equipment as above) were used to document them.

###### 1.3.2 Mandible morphology

To allow for inferences about the potential feeding ecology of the fossils, the MA was studied on a phylogenetically diverse sample of extant species including several orders of polyneopteran insects. Twenty-nine recent taxa of Polyneoptera (tab. S2) were investigated using micro-computed tomography (µCT) carried out at several synchrotron facilities: Beamline BW2 and IBL P05 of the outstation of the Helmholtz Zentrum Geesthacht at the Deutsches Elektronen Synchrotron (DESY), the beamline TOMCAT at the Paul Scherrer institute (PSI), the TOPO-TOMO beamline of the Karlsruhe Institute of Technology (KIT), and beamline BL47XU of the Super Photon Ring 8GeV (SPring-8).

##### 1.4 Analysis of the mandibular mechanical advantage

###### 1.4.1 Introduction

The mechanical advantage (MA) is a straightforward biomechanical metric which was first introduced for vertebrates (*59*) and was used since in studies on vertebrate and arthropod jaw mechanics (*52, 60, 61*). The MA is defined as the inlever to outlever ratio. For dicondylic insect mandibles the inlever is the distance between the application of the input force and the joint axis, while the outlever arm is the distance from the biting point to the joint axis (fig. S1).

The MA thus indicates the percentage of force transmitted to the food item (i.e. the effectivity of the lever system). Although more detailed investigations concerning muscular insertion angles, muscle volumes, spatial arrangements and muscle characteristics would be needed to quantify the absolute forces applied to a given food item, the MA is a useful mechanical performance index: It allows a size independent comparison of the relative efficiencies within the mandibular lever system and it can be readily measured in a wide array of dried museum specimens as well as freshly collected ones. Here, we used it to assess the efficiency of the mandibular lever system of insect fossils for the first time.

Automatic segmentations of the mandibles were performed using the software ITK-snap (*62*) after which STL files were imported into the software Blender (www.blender.org) for further processing (fig. S1). The gnathal edge was defined sensu Richter et al. (*63*) as the area from the pars molaris (proximal to the mouth opening) to the pars incisivus (distalmost tooth). Since the homology of subparts of the gnathal area is debated (*63-65*), the gnathal outline, as seen when orienting the mandible in line with the rotation axis (fig. S1), was scaled as a percentage of tooth row length. For this, ∼800 points for each specimen were wrapped against the gnathal outline in Blender and the distance between each point orthogonal to the mandibular rotation axis (= outlever) was measured. Similarily, one point was placed at the insertion point of M. craniomandibularis internus on the mandible and the distance between this point orthogonal to the rotation axis was measured (i.e. inlever). MA measurements were carried out on the segmentations of the left mandible for each specimen. All measurements and calculations were carried out in the R software environment (v. 1.1.383) using custom scripting. Separate MAs for each studied fossil were computed and combined to a composite fossil mandible representation (CFMR) using a Procrustes superimposition as implemented in geomorph v.3.0.5. in order to account for uncertainties in MA extraction due to potential distortion artefacts. From this superimposition the mean MA shape was extracted and used together with the MAs of recent species for the further analysis steps. Polynomial functions of the 1^st^ – 20^th^ order were fitted against all MA profiles. The Akaike information criterion (AIC) was used to determine the polynomial function with the best relative fit whose coefficients were then used for further analysis.

###### 1.4.2 Phylogenetic signal

Phylogenetic signal was assessed using the most recent phylogenetic estimate of the 1kite consortium (www.1kite.org) as a basis (*66*). The phylogeny was pruned in order to contain only the taxa analysed here. The fossils were fitted into the phylogenetic estimate based on inference derived from wing venation and leg and ovipositor morphology (Fig. 2). Specifically, *Ct. frequens* possesses a fusion of CuA (emerging from M+CuA) with CuPa, which is the defining character state of the taxon Archaeorthoptera (*67*), and **olis2**, the defining character state of the taxon Neoclavifera tax. n. (see below). Both taxa include crown-Orthoptera and some stem-relatives. The lack of a branching of CuPa (which would indicate a panorthopteran), and of jumping hind-legs (which would indicate a saltatorian) both indicate that *Ct. frequens* is not a crown-orthopteran.

Phylogenetic signal was assessed using the K statistic as implemented in geomorph v.3.0.5 (*54*) with 10,000 random permutations. This test statistic was found to be the most efficient approach to test for phylogenetic signal (*68*). Since significant phylogenetic signal was detected, a principal component analysis (PCA) as well as phylogenetic PCA as implemented in the phytools package v.0.6-44 (*55*) were carried out in order to compare the analysed specimens in MA shape space.

#### 2 Results

##### 2.1 Systematic Palaeontology

In this section the systematics at the family-group level and below conforms to the ICZN to ensure that the new species name is valid under this Code, while that above the family-group, left ungoverned by the corresponding code, conforms to the principles of cladotypic nomenclature (*69*; and subsequent accounts), itself compliant with the PhyloCode (*70*). Specifically, a cladotypic definition corresponds to an apomorphy-based definition using two species as internal specifiers (each being anchored to a specimen designated as type). There are minor discrepancies between cladotypic nomenclature practice on the one hand and recommendations of the PhyloCode on the other. Notably, the first author to have associated the selected defining character state and a taxon is to be acknowledged under the former procedure.

###### 2.1.1 Nomenclature above family-group level

Taxon *Neoclavifera* Béthoux, new clade name (*nom*. Béthoux *n*., *dis*. Kluge, 2016 (*14*), *typ*. Béthoux *n*.)

###### Registration number

Xxxx.

###### Definition

Species that evolved from the hypothetical ancestral species in which the character state ‘in female ovipositor, occurrence of a locking mechanism composed of a rhachis on gonostylus IX and of an aulax on gonapophysis IX’ (also called ‘secondary olistheter’; as opposed to ‘in female ovipositor, absence of a locking mechanism composed of a rhachis on gonostylus IX and of an aulax on gonapophysis IX’), as exhibited by *linderi* Dufour, 1861 (currently assigned to the genus *Dolichopoda* Bolívar, 1880) and *artinii* Griffini, 1913 (currently assigned to the genus *Homogryllacris* Liu, 2007), was acquired.

###### Abbreviated definition

∇ apo secondary olistheter [*Dolichopoda linderi* (Dufour, 1861) & *Homogryllacris artinii* (Griffini, 1911)].

###### Etymology

From ‘neo-’, Ancient Greek prefix for ‘new’; ‘clavis’, Latin for ‘key’; and ‘-fera’, Latin suffix for ‘bearing’ (feminine). This is a direct reference to **olis2**, which **rh** resembles a key in cross-section.

###### Reference phylogeny

The monophyly of Orthoptera, which includes all extant species sharing the defining character state of *Neoclavifera*, is beyond doubt. Wipfler *et al*. (*71*) and Song *et al*. (*66*) compose two recent accounts on the topic. It follows that the acquisition of the defining character state in the cladotypic species / specifiers, attested by Cappe de Baillon (*12*), very probably occurred once (*14*). It is considered lost in Caelifera.

###### Qualifying clauses

Several qualifying clauses are explicit when using a cladotypic definition, but they need to be specified for a PhyloCode usage. The name *Neoclavifera* shall be considered as invalid as that of a taxon if it occurs that (i) the defining character state was acquired by the cladotypes / specifiers convergently, (ii) the defining character state is a plesiomorphy, (iii) the cladotypes / specifiers belong to a single species and/or (iv) the defining character state does not occur in the specifiers (unless it is secondarily lost). There is no known evidence that one of these clauses might be challenged in our case.

###### Composition

All Saltatoria (including all extant Orthoptera) and ‘lobeattid insects’ (understood as including cnemidolestodeans) (and see section 2.1.2, taxonomic discussion).

###### Discussion

At the first glance, the name Ensifera Chopard, 1920, appears as a suitable name to convert. However, it is an explicit reference to the sword-like shape of the ovipositor valves in the corresponding insects, which composes a pre-occupation under cladotypic nomenclature (conversely, the taxon name Caelifera Ander, 1936, is an explicit reference to chisel-like shape of the valves). In other words, the name etymologically refers to a character state different from that used to define the new taxon, which makes it unavailable for the aimed purpose. The same applies to the taxon name Dolichocera Bei-Bienko, 1964 (‘long horned’; and, conversely, ‘Brachycera Bei-Bienko, 1964’ for ‘short horned’), favoured by Kluge (*14*). Moreover, current classificatory schemes customarily regard Ensifera and Caelifera as sister-groups, while our results predict that Caelifera is to be nested within Ensifera. Prolonged ambiguity on the conversion of ‘Ensifera’ as a defined taxon is then to be expected, not mentioning the fact that *Ensifera* Lesson, 1843 is a genus name for sword-billed hummingbirds, and *Ensifera ensifera* (Boissonneau, 1839) its type species.

Given this situation, and the absence of a name composing a direct reference to the occurrence of **olis2**, we propose to coin a new one accordingly. Based on our literature survey, Kluge (*14*) is the first author to have discriminated a taxon on the basis of the defining character state only. This author stated that an **olis2** is the autapomorphy of ‘Dolichocera’, but the name being a direct reference to another character state (see above), it follows that a new one is needed, hence *Neoclavifera*.

The meaning of the terms ‘rhachis’ and ‘aulax’ is critical to the proposed definition. Modest elevation and groove likely made the transition from adjoined smooth surfaces to ones bearing a proper rhachis and aulax. It is therefore necessary to define rhachis and aulax, as follows: a rhachis is a projection whose base is narrower than its projected part at its widest (best assessed in cross-section), and an aulax is its counterpart.

As defined, and based on species currently known, the composition of the taxa *Archaeorthoptera* and *Neoclavifera* overlap. We hypothesize that the defining character state of *Archaeorthoptera* was acquired in a hypothetical ancestral species distinct from the one of *Neoclavifera*, but the order of acquisition of their respective defining character states remains unknown.

###### 2.1.2 Nomenclature at the family-group level and below

Family Ctenoptilidae Aristov, 2014

Genus *Ctenoptilus* Lameere, 1917

*Ctenoptilus frequens* Chen, Gu & Béthoux, sp. nov.

###### Etymology

Based on ‘*frequens*’ (‘frequent’ in Latin), referring to the abundance of the species at Xiaheyan.

###### Holotype

Holotype: CNU-NX1-326 (female individual; Fig. 1).

###### Referred material

CNU-NX1-198, −199, −731 to −759, −764 (specimens herein figured: CNU-NX1-198, −199, −731, −732, −738, −740–744, −747–754, 756–759).

###### Locality and horizon

Xiaheyan Village, Zhongwei City, Yanghugou Formation (Ningxia Hui Autonomous Region, China); latest Bashkirian (latest Duckmantian) to middle Moscovian (Bolsovian), early Pennsylvanian (*4*).

###### Differential diagnosis

Compared with *Ctenoptilus elongatus* (Brongniart, 1893), its most closely related species (*SI Appendix*, section 2.1), smaller size (deduced from forewing length) and prothorax longer than wide (as opposed to quadrangular).

###### General description

Body length (excluding antennae, including ovipositor) about 42–52 mm (based on female individuals only). **Head**: prognathous, head capsule heart-shaped in dorsal view; **md** with strongly sclerotized and prominent incisivi and a well-sclerotized molar area; **la** with a strong apical tooth and a smaller sub-apical one; **mp** well-developed, with 5 observed segments; tentorium composed of well-developed **ata, ct** and **pta**, dorsal arms not visible; **co** located in the midline along the dorsal side of the head capsule, then branching into two diverging **fc**; **ant** long, filiform. **Thorax**: prothorax longer than wide, longer than head; boundary between mesothorax and metathorax not visible. *Wings:* ScP reaching RA distal to the two thirds of wing length; RA with few or no anterior veinlets; RA and RP strong, parallel for a long distance; RA-RP area narrow in its basal half; at the wing base, R and M+CuA distinct; MA and MP simple for a long distance, with similar numbers of terminal branches, usually 1–3, rarely more than 4; CuA diverging from M+CuA and fusing with CuPa; CuA+CuPa posteriorly pectinate. *Forewing*: length 31.5/**36.1**/41.2 mm, largest width 6.9/**8.3**/10.7 mm, membranous; ScP with anterior veinlets; RA-RP fork slightly distal to the point of divergence of M and CuA (from M+CuA); RP branched distally, near the second third of wing length, usually with 11–17 branches reaching apex, and occasionally 1–2 veinlets reaching RA; first split of M+CuA (into M and CuA) near the first fourth of wing length; between the origin of CuA (from M+CuA) and the first fork of RP, M very weak; first fork of M near wing mid-length; MA distinct from RP, connected to it by a short cross-vein, or occasionally fused with it for a short distance; median furrow located along M and then MP; CuA+CuPa with most of its main branches further branched, with a total of 16–26 terminal branches; in basal part, CuA+CuPa emitting strong posterior veinlets, vanishing before they reach the claval furrow; CuPb concave, weak and simple; AA1 with 3–4 branches; AA2 with about 10 branches; cross-veins mostly not reticulated, except along the apical and postero-apical section of the wing margin, and in the ScP/ScP+RA-RP area (where they are particularly strong); longitudinal pigmented areas located (i) along R, (ii) along CuA, and then the main stem of CuA+CuPa and (iii) along the posterior wing margin, distal to the endings of the first branches of CuA+CuPa; these three areas merge distally; additional pigmented area along AA1. *Hind wing*: as in forewing, except for the following: slightly shorter than forewing; RA-RP fork opposite the point of divergence of M and CuA (from M+CuA); RP usually with 11–16 branches reaching apex; M forked at the first quarter of the wing; M with 5–8 branches reaching posterior wing margin; CuA+CuPa with 5–8 branches; pigmented area forming an arc covering the apex, beginning along RA and ending close to the end of CuPb; plicatum well developed, with plica prima anterior reaching the posterior wing margin opposite the end of ScP (on RA). *Legs*: Fore-leg femur 4.9–6.3 mm long, 1.0–1.3 mm wide, tibia 5.2–6.3 mm long; mid-leg femur 5.2–6.4 mm long, tibiae 5.9–7.3 mm long; hind-leg femur 7.5–11.5 mm long, tibia 9.8–12.0 mm long; spines, probably in two rows, present along the ventral side of tibia of all legs, concentrated near the apex (fore-leg, at least 12 spines; mid-leg, at least 8 spines; hind-leg, at least 15 spines); tarsus 5-segmented, 2^nd^, 3^rd^ and 4^th^ segments shorter, terminal tarsal segment with paired claws and arolium (deduced from well-preserved fore-legs). **Abdomen**: abdomen about 17–23 mm long (based on female individuals only); female with a prominent sword-like ovipositor (see more detailed interpretation below and specimens description).

###### Specimens description. Holotype, CNU-NX1-326 (Fig. 1)

Positive and negative imprints of an almost complete female individual, viewed dorsally, very well preserved, with head, thorax, leg remains (including well exposed fore-legs) and complete right forewing; right hind apex missing, left wings incomplete, left hind wing very incomplete, ovipositor apex concealed under right forewing. **Head**: about 6.6 mm long, 4.3 mm wide, prognathous; mandibles about 2.0 mm long, with prominent teeth at their apex; gnathal edge of right **md** clearly visible, heavily sclerotized, with the distal incisivus shorter than the subdistal ones; **mp** strong, but segments not visible; **f** large and separated to the vertex by a U-shaped line, laterally delimited to the well-developed genal area by a line; frontal and coronal sutures well-developed, located at the closest distance of the eyes to each other; Eyes large, laterally protruding from the head capsule covering about half of the lateral head profile; **ant** incomplete, 6.3 mm long as preserved. **Thorax**: prothorax about 5.5 mm long, 3.7 mm wide. *Left forewing*: preserved length 22.2 mm, best width 8.1 mm; M with its 2 main branches preserved, CuA+CuPa with 22 terminal branches preserved. *Right forewing*: length 32.6 mm, width 9.6 mm; RP simple for 14.3 mm, with 16 branches reaching wing apex and 1 reaching ScP+RA; MA connected to RP by a short cross-vein, with 3 branches, MP with 4 branches; CuA+CuPa with 26 terminal branches preserved; CuPa partly preserved. *Right hind wing*: preserved length: 30.4 mm, best width 8.6 mm; plicatum creased. *Legs*: fore-leg femur about 4.9 mm long and 1.2 mm wide, tibia 6.3 mm long and 0.7 mm wide, tarsus about 5.0 mm long, tarsal segments (5), paired claws and arolium visible; mid- and hind-legs incomplete and/or not well exposed. *Legs*: spines well exposed on foreleg tibiae and distal part of a mid-leg tibia. **Abdomen**: bent (probably a consequence of decay), about 17 mm long, ovipositor viewed laterally, possibly slightly obliquely; bases of **gp8** strongly sclerotized, well visible.

###### CNU-NX1-749 (Fig. 2A and B, and fig. S7A–E)

Positive and negative imprints of an almost complete female individual, wings incomplete and overlapping, body about 45 mm long. **Head:** about 6.4 mm long, 3.5 mm wide. **Thorax:** prothorax about 5.6 mm long, 3.7 mm wide. *Legs*: fore-leg femur 4.9 mm long, 1.2 mm broad, tibia 5.8 mm long, 0.8 mm broad, tarsus about 3.8 mm long; mid-leg femur 5.9 mm long, 1.0 mm broad, tibiae 7.3 mm long, 0.8 mm broad, tarsus about 4.9 mm long; hind-leg femur 6.1 mm long, 1.1 mm broad, tibia 10.1 mm long, 0.7 mm broad; spines visible, or even well-exposed, on each exposed tibiae. **Abdomen**: about 17 mm long (excluding ovipositor); sword-like ovipositor viewed laterally, about 8.4 mm long; antero-basal apophyses of **gs9, gp9** and **gp8** distinct, well delineated; near the ovipositor base, dorsal and ventral edges of **gs9** and **gp8**, and ventral edge of **gp9** well delineated; dorsal edge of **gp9** visible in the distal half of the ovipositor; **olis1** and **olis2** visible near the ovipositor base, strongly sclerotized; **olis1** located along the ventral edge of **gp9** and dorsal edge of **gp8**; **olis2** located close to (or along) the ventral edge of **gs9**, and laterally on **gp9**; **olis1** and **olis2** converging; ventral edge of **gp8** with teeth more prominent and densely distributed near the apex.

###### CNU-NX1-742 (Fig. 2C and D, and fig. S8A–C)

Positive and negative imprints of an almost complete female individual, partly disarticulated, left forewing missing; body about 52 mm long. **Head**: detached from the rest of the body, mouthparts not discernible. **Thorax**: prothorax about 7.0 mm long, 3.6 mm width. *Wings*: a forewing and two hind wings visible, poorly preserved. *Legs*: fore-leg femur 5.7 mm long, 1.1 mm broad, tibia 6.2 mm long, 0.9 mm broad; spines well exposed on one hind-leg tibia, some visible on one fore-leg tibia. **Abdomen**: strongly bent, segments not discernible; ovipositor very well preserved, detached from the rest of the abdomen, about 9.5 mm long; antero-basal apophyses of **gs9, gp9** and **gp8** distinct, well delineated; near the ovipositor base, dorsal and ventral edges of **gs9** and **gp8**, and ventral edge of **gp9** well delineated; dorsal edge of **gp9** visible at the extreme base and in the distal half of the ovipositor; **olis1** and **olis2** visible near the ovipositor base, strongly sclerotized; **olis1** located along the ventral edge of **gp9** and dorsal edge of **gp8**; **olis2** located close to (or along) the ventral edge of **gs9**, and laterally on **gp9**; **olis1** and **olis2** converging; ventral edge of **gp8** with teeth more prominent and densely distributed near the apex.

###### CNU-NX1-754 (Fig. 3A and B, and fig. S7I–K)

Positive and negative imprints of an almost complete individual, well-preserved, wings overlapping, incomplete and partly creased, end of abdomen missing. **Head**: about 6.8 mm long 4.5 mm wide; **md** with strongly sclerotized and prominent incisivi and a well-sclerotized molar area; terminal teeth of **la** visible; **ca** distinguishable; **co** located in the midline along the dorsal side of the head capsule, then branching into two diverging **fc. Thorax**: prothorax about 5.9 mm long, 4.4 mm wide. *Legs*: fore-leg femora 5.3 mm long and 1.1 mm broad, tibiae 5.2 mm long and 0.7 mm broad, tarsus about 4.0 mm long; mid-leg femur 5.4 mm long and 1.1 mm broad, tibia 5.9 mm long and 0.8 mm broad, tarsus about 4.5 mm long; fore- and mid-leg tarsi well preserved, 5-segmented with paired claws and arolium; 2^nd^, 3^rd^ and 4^th^ segments shorter, ventral process (projecting forward) of 3^rd^ and 4^th^ segments visible; hind-leg femora 7.5 mm long; end of hind-leg tibiae missing, 7.1/5.9 mm long, 0.7 mm broad; spines well-exposed on one of the forelegs tibiae. **Abdomen**: about 14 mm as preserved, segments not discernible.

###### CNU-NX1-764 (Fig. 3C and D)

Positive and negative imprints of an almost complete, isolated head, posterior part possibly overlapping with prothorax; mouthparts well preserved; **md** in occlusion, 2.1 mm long, 1.1 mm wide at their base, provided with strongly sclerotized and prominent incisivi and a well-sclerotized molar area; distal part of **la** visible, provided with a strong apical teeth and a smaller sub-apical one; tentorium composed of well-developed **ata, ct** and **pta**, dorsal arms not visible; **ct** 1.2 mm long and 0.3 mm wide.

###### CNU-NX1-752 (fig. S2A and B)

Positive and negative imprints of a partly incomplete individual, head and prothorax well exposed, a single fore-leg preserved, wings partly spread, right hind wing creased, most of abdomen missing. **Thorax:** prothorax about 7.0 mm long, 4.0 mm wide; *Right forewing*: preserved length 35.1 mm, width 8.8 mm. RP simple for 14.9 mm, with 12 branches preserved; M poorly preserved, MA simple, MP with 3 branches reaching the posterior wing margin; CuA+CuPa incomplete, with 15 visible branches. *Left forewing*: apex missing, preserved length 32.3 mm, width 8.7 mm; M not visible in its median portion; a portion of CuPa basal to its fusion with CuA visible. *Left hind wing*: length 29.5 mm, width 10.0 mm; plicatum in resting position and creased; RP with 11 branches reaching apex; M with 8 terminal branches.

###### CNU-NX1-738 (fig. S2C and D)

Imprint of an individual with parts of prothorax and thorax preserved, a left forewing (as negative imprint) and a right hind wing (as positive imprint). *Left forewing*: length 31.3 mm, width 10.3 mm; RP simple for 15.2 mm, with 13 branches reaching wing apex; MA connected to RP by a very short cross-vein; M with a total 5 distal branches (MP simple); CuA+CuPa with 15 preserved terminal branches. *Right hind wing*: partly creased, plicatum not discernible/preserved, wing base not discernible; length 29.6 mm, width 7.2 mm; RP with 13 branches reaching wing apex.

###### CNU-NX1-759 (fig. S3A and B)

Imprint of a nearly complete individual, most of head missing, left forewing twisted, right hind wing concealed under right forewing, a mass of circular cavities probably indicates the location of abdominal remains. **Thorax:** prothorax about 6.4 mm long, 3.1 mm wide; *Right forewing*: apex missing, anal area not discernible; length 34.1 mm, best width 8.0 mm; RP simple for 13.7 mm, with 11 branches preserved; one reaching ScP+RA; MA fused with RP for 0.6 mm, with 2 terminal branches, MP with 2 branches; CuA+CuPa not fully discernible, with 16 terminal branches preserved. *Right hind wing*: anterior wing margin and plicatum not discernible; RP with 7 branches preserved; M with 5 branches (2,3), CuA+CuPa incomplete, with 4 branches. *Legs*: left legs almost missing, right fore- and mid-leg with femur and tibia preserved; right fore-leg, femur 5.7 mm long, tibia 5.3 mm long; right mid-leg, femur 6.4 mm long, tibia 8.4 mm long; right hind leg, femur 8.3 mm long, tibia 11.1 mm long, tarsus about 6.6 mm long, with 5 tarsal segments, claws and arolium visible; spines visible on one of the hind-leg tibiae.

###### CNU-NX1-750 (fig. S3C and D)

Positive and negative imprints of an almost complete individual, forewings overlapping hind wings, complete set of legs, abdomen poorly preserved and incomplete. **Thorax**: prothorax about 5.9 mm long, 4.2 mm wide. *Right forewing*: preserved length 32.2 mm, best width 8.6 mm; RP simple for 14.2 mm, with 11 branches preserved; MA with 2 branches, MP with 3 branches; CuA+CuPa with 23 terminal branches, CuPb partly visible. *Left forewing*: apex missing, posterior wing margin not discernible; RP simple for 13.49 mm, CuA+CuPa with 18 branches preserved. *Hind wings*: apices and most of margins missing/not discernible, plicata partly unfolded, creased. *Right hind wing*: preserved length 28.3 mm; M with 10 branches reaching apex, CuA+CuPa with 6 branches preserved. *Left hind wing*: basal part not discernible; preserved length 22.7 mm, width 8.7mm; RP with 4 branches preserved, CuA+CuPa with 5 branches preserved. *Legs*: fore-leg poorly preserved; mid-leg femur 5.4 mm long, 1.1 mm broad, tibia 6.7 mm long, 0.8 mm broad; hind-leg femur 7.9 mm long, 1.3 mm broad, tibia 12.3 mm long, 0.8 mm broad.

###### CNU-NX1-731 (fig. S3E and F)

Positive and negative imprints of an almost complete individual, very well-preserved, legs and left hind wing missing; abdomen broken. **Head**: preserved length 6.7 mm. **Thorax**: prothorax about 4.7 mm long, 3.5 mm wide. *Left forewing*: length 39.6 mm, best width 9.1 mm; RP simple for 13.4 mm, with 10 branches reaching apex and one branch reaching with ScP+RA; M well preserved, connect with RP by a long, oblique cross-vein; MA with 3 branches, MP with 1 branch; CuA+CuPa with 17 branches reaching the posterior wing margin, CuPb poorly preserved. *Right forewing*: preserved length 35.9 mm, best width 9.2 mm; a vein interpretable as ScA partly preserved; RP simple for 13.1 mm, with 10 branches reaching apex and one veinlet reaching with ScP+RA; M well preserved, connected with RP by a long, oblique cross-vein; MA with 2 branches, MP with 2 branches; CuA+CuPa with 19 branches reaching the posterior wing margin; AA area with serval branches preserved. *Right hind wing*: apex missing; RP with 10 branches directed towards apex and 2 veinlets reaching with ScP+RA; MA and MP simple for a long distance, with 3 and 2 branches reaching the posterior wing margin, respectively; CuA+CuPa with 6 terminal branches preserved; plicatum folded, creased.

###### CNU-NX1-747 (fig. S4A–D)

Positive and negative imprints of an almost complete individual, left forewing and right hind leg missing. **Head**: about 6.0 mm long, 4.4 mm wide; **md** about 1.7/2.0 mm long, 1.4 mm wide at their base; apical tip of **la** visible, **ct** 0.9 mm long 0.2 mm wide; compound eye oval; circumoccular ridge well developed. **Thorax**: prothorax about 5.5 mm long, 3.3 mm wide. *Right forewing*: preserved length about 29 mm, best width 8.2 mm; RP simple for 12.3 mm, with 10 branches, two of them reaching ScP+RA; MA and MP with two branches each; CuA+CuPa with 19 terminal branches visible. *Hind wings:* plicatum folded, with numerous anal veins, not clearly discernible. *Left hind wing*: length 30.1 mm, best width 7.8 mm; RP simple for 9.6 mm, with 11 branches reaching wing apex and a single veinlet reaching ScP+RA; MA and MP simple for a long distance, each with 3 branches; CuA+CuPa posteriorly pectinate, with 6 terminal branches. *Right hind wing*: overlapping with right forewing, only partly discernible; RP simple for 8.3 mm, with 9 branches preserved. *Legs*: right legs poorly preserved and/or incomplete; left fore-leg femur 6.3 mm long and 1.0 mm wide, tibia 5.2 mm long and 0.6 mm wide; mid-leg femur 5.0 mm long and 1.1 mm wide, tibia 6.8 mm long and 0.6 mm wide; hind-leg femur 8.6 mm and 1.0 mm wide, tibia 9.8 mm long and 0.6 mm wide; spines well exposed on both foreleg tibiae, and the preserved mid-leg and hind-leg tibiae.

###### CNU-NX1-741 (fig. S4E and F)

Positive and negative imprints of an incomplete individual, left forewing and thorax preserved, left hind wing and right forewing incomplete. *Left forewing*: apex missing, preserved length 34.8 mm, best width 10.7 mm; RP simple for 15.6 mm, with 7 branches preserved; M poorly preserved, MA and MP with 3 branches each; CuA+CuPa with 23 terminal branches; AA1 with 4 main branches. *Right forewing*: only the basal part preserved, AA1 with 4 branches, AA2 with 6 branches preserved. *Left hind wing*: preserved length 33.1 mm; RP with 6 branches; MA and MP simple in the preserved part.

###### CNU-NX1-748 (fig. S5A and B)

Well preserved isolated right forewing, negative imprint; length 38.4 mm, best width 8.7 mm; RP simple for 14.1 mm, with 12 terminal branches reaching apex; MA with 2 branches, the anterior one connected to RP by a cross-vein; MP with 5 branches; CuA+CuPa with 16 terminal branches reaching the posterior wing margin, and a branch fused with MP; CuPb visible, area between CuPb and AA1 narrow; AA1 with 4 preserved branches.

###### CNU-NX1-732 (fig. S5C and D)

Positive imprint of nearly complete right forewing; preserved length 30.3 mm, best width 6.6 mm; RP simple for 12.8 mm, with 17 branches reaching wing apex and 3 branch reaching ScP+RA; basal portion of M relatively well-preserved, first fork located opposite wing mid-length; MA simple; MP with 3 branches reaching posterior wing margin and a veinlet fusing with MA; CuA+CuPa with 16 branches; AA1 with 3 branches.

###### CNU-NX1-757 (fig. S5E and F)

Negative imprint of a well-preserved, isolated right forewing; length 35.3 mm, best width 8.4 mm; RP simple for 14.4 mm, with 12 branches reaching wing apex and 2 branches reaching ScP+RA; basal portion of M relatively well-preserved; MA and MP with 4 and 3 branches, respectively; CuA+CuPa with 19 terminal branches reaching posterior wing margin; AA1 with 4 branches; AA2 poorly preserved.

###### CNU-NX1-758 (fig. S5G–H)

Negative imprint of a well-preserved, isolated left forewing, slightly creased along the claval furrow; length 36.6 mm, best width 7.2 mm; RP simple for 13.9 mm, with 11 terminal branches reaching apex; MA and MP with 2 and 3 branches, respectively; CuA+CuPa with 16 terminal branches reaching posterior wing margin. **CNU-NX1-744 (fig. S5I and J):** Negative imprint of a well-preserved, isolated right forewing; length 41.2 mm, best width 8.7 mm; RP simple for 16.0 mm, with 12 terminal branches visible; MA connected to RP by a short cross-vein; MA and MP with 2 terminal branches each; CuA+CuPa with 19 branches reaching posterior wing margin; AA1 with 3 preserved branches.

###### CNU-NX1-751 (fig. S5K and L)

Negative imprint of nearly complete forewing pair and apical fragment of a hind wing. *Forewings*: basal half of M and CuPb not visible; MA and MP with 2 branches each. *Left forewing*: preserved length 30.4 mm, best width 6.9 mm; RP simple for 13.7 mm, with 17 branches reaching wing apex; CuA+CuPa with 17 terminal branches, AA1 with 3 branches. *Right forewing*: length 27.1 mm, best width 7.4 mm; RP simple for 13.6 mm, with 14 preserved branches reaching wing apex; CuA+CuPa with 21 terminal branches.

###### CNU-NX1-743 (fig. S5M–Q)

Positive imprint of an incomplete right forewing and of an ovipositor. *Right forewing*: basal part missing, preserved length 26.9 mm, best width 7.8 mm; RP with 14 branches reaching wing apex and a veinlet reaching ScP+RA; M and most of MA poorly preserved; MA and MP with 3 branches each; CuA+CuPa incomplete, with 17 branches reaching the posterior wing margin, and 1 veinlet fusing with MP. *Ovipositor*: preserved length 9.2 mm; **olis2** and dorsal margin of **gp8** strongly sclerotized; prominent teeth visible in the distal part of **gp8**.

###### CNU-NX1-198 (fig. S6A–B)

Positive and negative imprints of an almost complete individual, head and abdomen missing, wings moderately well preserved, left wings overlapping. **Thorax**: prothorax about 6.6 mm long, 3.9 mm wide. *Right forewing*: length 32.1 mm, best width 8.8 mm; RP simple for 12.1 mm, with 9 branches preserved; M poorly preserved, MA and MP with 3 and 2 branches, respectively; CuA+CuPa incomplete, with 14 branches preserved. *Right hind wing*: length 28.7 mm, best width 14.6 mm; RP simple for 14.6 mm, with 8 branches preserved; MA and MP simple for a long distance, M with 7 branches reaching the posterior wing margin; fusion of CuA (emerging from M+CuA) with CuPa visible; CuA+CuPa with 7 terminal branches; CuPb partly preserved; plicatum almost fully deployed, large, probably with vannal folds; AA with 9 branches preserved.

###### CNU-NX1-740 (fig. S6C and D)

Positive and negative imprints of an incomplete individual, with forewings and right hind wing poorly preserved, abdomen not discernible. **Head**: 7.5 mm long, 4.7 mm wide. **Thorax**: prothorax about 5.5 mm long, 4.5 mm wide. *Left hind wing*: apex missing; fusion of CuA (emerging from M+CuA) with CuPa visible; plicatum well deployed, large, with several veins preserved (attributable to AA). *Legs*: fore-leg femur length 5.5 mm long and 1.2 mm wide; mid-leg femur 5.2 mm long and 1.2 mm wide; hind-leg femur 11.5 mm long and 1.2 mm wide, tibiae 12.0 mm long and 0.8 mm wide, tarsus about 6.2 mm long, paired claws and arolium preserved.

###### CNU-NX1-199 (fig. S6E and F)

Positive and negative imprints of isolated right hind wing; wing base not discernible, apex missing; preserved length 17.4 mm, best width 10.1 mm; RP simple for 8.9 mm, with 6 branches preserved; M with 5 branches reaching the posterior wing margin; CuA+CuPa with 8 terminal branches; plicatum well-deployed, with 17 branches preserved (attributable to AA).

###### CNU-NX1-753 (fig. S6G and H)

Negative imprint of an isolated left hind wing, plicatum not discernible/preserved; length 34.2 mm, best width 10.1 mm; at the wing base, M+CuA distinct from R; RP simple for 9.5 mm, with 16 branches reaching wing apex; MA and MP simple for a long distance, with 3 and 4 branches, respectively; CuA+CuPa with 5 terminal branches preserved; plicatum with several visible veins (attributable to AA).

###### CNU-NX1-756 (fig. S7F–H)

Positive and negative imprints of an almost complete female individual, wings poorly preserved and incomplete, total length (excluding **ant**) about 51 mm. **Head**: 7.1 mm long, 3.6 mm wide; **md** open; left **md** with well-discernible **il** and **mo**; left **la** with a strong apical tooth and a smaller sub-apical one; **co** located in the midline along the dorsal side of the head capsule, then branching into two diverging **fc**; **ant** long, filiform; **ce** 1.4 mm long and 0.8 mm wide. **Thorax**: prothorax about 5.8 mm long, 4.0 mm wide. **Abdomen**: length about 23 mm, segments not discernible; exposed portion of ovipositor about 5.0 mm long.

###### Taxonomic discussion

The new species is closely related to a number of Pennsylvanian insects collectively referred to as ‘lobeattids’ and characterized by (i) a RA/RP fork located basally, (ii) a RA-RP area widening sharply distal to the end of ScP (on RA) and (iii) CuA+CuPa with one main anterior branch posteriorly pectinate and with abundant branches (commonly, ca. 20) reaching the posterior wing margin. This assemblage includes *Eoblatta robusta* (Brongniart, 1893) and *Ctenoptilus elongatus* (Brongniart, 1893), from the Commentry locality (France); *Lobeatta schneideri* Béthoux, 2005, *Anegertus cubitalis* Handlirsch, 1911 and *Nectoptilus mazonus* Béthoux, 2005, from Mazon Creek (USA); *Nosipteron niedermoschelensis* Béthoux & Poschmann, 2009, from Niedermoschel (Germany); *Lomovatka udovichenkovi* Aristov, 2015, from Lomovatka (Ukraine); *Beloatta duquesni* Nel, Garrouste & Roques; and *Sinopteron huangheense* Prokop & Ren, 2007, *Chenxiella liuae* Liu, Ren & Prokop, 2009, *Longzhua loculata* Gu, Béthoux & Ren, 2011 and *Protomiamia yangi* Du, Béthoux, Gu & Ren, 2017 from Xiaheyan (China). The genus *Miamia* Dana, 1864 and the order Cnemidolestodea are derived members of this assemblage.

Compared with known species, the new one is mostly similar to *Ct. elongatus, Ne. mazonus* and *Lom. udovichenkovi* owing to the elongate to very elongate shape of the forewing (presumed in the latter). A further similarity of the new species with *Ct. elongatus* and *Lom. udovichenkovi* is the occurrence of numerous posterior basal veinlets of CuA+CuPa vanishing before reaching CuPb. Strikingly, the new species and *Ct. elongatus* share a very particular forewing coloration pattern, with three longitudinally-orientated, pigmented bands. We therefore propose to assign the new species to the genus *Ctenoptilus* Lameere, 1917.

Note that Béthoux & Nel (*1*) identified, in one specimen of *Ct. elongatus*, a linear structure they interpreted as MP, that would indicate a basal position of the first fork of M. However, based on data on the new species and on the original descriptions of *Ne. mazonus* and *Lom. udovichenkovi*, we assume that the ‘linear structure’ is more likely the median furrow alone. If so, the first fork of M, in *Ct. elongatus*, might well be located closer to the middle of the forewing, as in the new species and in *Ne. mazonus* and *Lom. udovichenkovi*. Note that this fork is located more basally in *Lom. udovichenkovi* than in the new species.

The forewing of the new species is smaller than in *C. elongatus*. Even though post-depositional deformation is known to have occurred at Xiaheyan and might have artificially elongated the forewing, the longest forewing of the new species is ca. 40 mm (with an average at ca. 36 mm), while *Ct. elongatus* forewings are 45–50 mm long. Note that female-biased sexual size dimorphism is known in a related species of Pennsylvanian Archaeorthoptera (*72*). However, if one assumes that all know specimens of *Ct. elongatus* are males, then females of this species would be even longer. If all know specimens of *Ct. elongatus* are females, then they can be compared with the longest representatives of the new species, but the size gap remains then. Finally, it remains possible that the difference in size is due to a latitudinal gradient (with *Ct. elongatus* living in the equatorial area, the new species at higher latitude), but available data on the impact of latitude in extant insects size variation is too contentious to provide matter for a grounded comparison (*73*). In summary, differences in size were considered sufficient to erect a new species.

Several specimens of the new species display a prothorax longer than wide (figs S1A and B, S2, and S6A,I), while it is more quadrangular in *Ct. elongatus* (*74*). It should be acknowledged, however, that the proportions of the prothorax in the holotype (Fig. 1) are similar to those of *Ct. elongatus*.

The set of specimens we investigated all share the coloration pattern typical for both *Ct. elongatus* and *Ct. frequens*. However, they display some variation in the forewing venation. The set of specimens on one hand, and, on the other, data on a few related species for which intra-specific variability was documented, demonstrate that this variability falls within the range of intra-specific variation. Lobeattid species relevant for comparison are *Lon. loculata, Miamia bronsoni* Dana, 1864 and *Miamia maimai* Béthoux, Gu, Yue & Ren, 2012.

Several specimens preserving a pair of sub-complete forewings (Fig. 1, and figs. S1A and B, S2C–F, and S4K and L) demonstrates that variation in the number and branching pattern of RP, M, and CuA+CuPa occur at the intra-specific level. More important variations are (i) the connection, or lack thereof, of an anterior veinlet from RP with RA, (ii) the connection, or lack thereof, of an anterior branch of MA with RP, and (iii) the connection of an anterior branch of CuA+CuPa with MP. As for (i), the set of specimens covers the complete range of variation, suggesting that it is not a character suitable to delimit species. Moreover, a similar range of variation has already been documented in *Lon. loculata* and *Miamia* spp. As for (ii), again, the set of specimens covers the complete range of variation of the character, with ‘an anterior branch of MA and RP distinct’ (figs. S3C and D, and S5K and L), ‘an anterior branch of MA and RP connected by a short cross-vein’ (Fig. 1, and figs. S2C and D, and S5A and B), ‘an anterior branch of MA and RP briefly connected’ (fig. S5I and J), and ‘an anterior branch of MA and RP fused for some distance’ (fig. S3A and B). Again, the same range of variation has been documented in *Lon. loculata* and *M. maimai*. As for (iii), the trait is very rare (fig. S4A and B). Given the above and the variation documented in *Lon. loculata*, it is of very minor relevance. In summary, observed differences in forewing venation are not sufficient to distinguish distinct species.

We assign several isolated hind wings, specifically the specimens CNU-NX1-199 (fig. S6E and F) and CNU-NX1-753 (fig. S6G and H) to *Ct. frequens* because they share the same size and the distinctive coloration of *Ct. frequens* hind wings, as documented from the holotype (Fig. 1) and other specimens preserving both fore- and hind wing, specifically CNU-NX1-752 (fig. S2A and B), CNU-NX1-738 (fig. S2C and D), CNU-NX1-731 (fig. S3E and F), CNU-NX1-747 (fig. S4A and B) and CNU-NX1-198 (fig. S6A and B). The specimen CNU-NX1-764 (Fig. 3C and D) is an isolated head. Compared with other species occurring at Xiaheyan, it can be confidently assigned to *Ct. frequens* based on its size, shape, and features of the mandibles. The specimens CNU-NX1-749 (fig. S6A–E), CNU-NX1-756 (fig. S6F–H), CNU-NX1-754 (fig. S6I–K) and CNU-NX1-742 (fig. S7A–C) can be confidently assigned to *Ct. frequens* based on size, wing venation and coloration, rectangular prothorax and/or long ovipositor.

##### 2.2 Ovipositor comparative analysis

This section complements schematic reconstructions provided in Fig. 2. Schemes representative of Grylloidea, Gryllacrididae and Anostostomatidae were derived from previous accounts (*12-14*).

###### 2.2.1 *Grylloblatta chandleri* Kamp, 1963 (schematized under ‘Grylloblattodea’ in Fig. 2G)

Our observations corroborate previous accounts (*15, 75*), in particular regarding the occurrence of a long **olis1** connecting **gp9** and **gp8**. Its **rh** is slightly dejected externally. We also noticed the occurrence of an olistheter interlocking left and right **gp9** along their dorsal margins. A specimen we observed had an egg engaged in the ovipositor. Due to the large diameter of the egg **olis1** unlocked, as well as the dorsal **gp9**–**gp9** olistheter. It can then be assumed that olistheters are comparatively labile structures in the species. In resting position (i.e. without engaged egg), when viewed externally, the ventral part of **gp9** is not concealed by **gp9**. Most of the area of **gp9** concealed by **gs9** is not as strongly sclerotized as its ventral part, except for the very base and its dorsal, ventral and apical margins.

###### 2.2.2 *Anacridium aegyptium* (Linnaeus, 1764) (schematized under ‘Caelifera’ in Fig. 2G)

Our observations corroborate previous accounts on other caeliferan species reporting the occurrence of an **olis1** connecting **gp9** and **gp8** along the entire ventral edge of the former (*14, 16*). Unlike reported by Ander (*57*), we found no evidence of an olistheter interlocking the ‘inner’ (i.e. **gp9**) and ‘posterior’ (i.e. **gs9**) valves (i.e. **olis2**). The **gp8** and Ander’s (*57*) ‘lateral basivalvular sclerite’ are extensively fused: they share the same lumen, and the dorsal and ventral fusion points are conspicuous in cross-section, owing to a clear invagination, coupled to a substantial and well-delimited thickening, of their shared wall.

###### 2.2.3 *Ceuthophilus* sp. (Fig. 2E and F; schematized under ‘Rhaphidophoridae’ in Fig. 2G)

We concur with previous accounts reporting that **olis2** occurs in this genus and in other Rhaphidophoridae (*12, 14, 76*). Unlike other orthopterans, the **rh** of **olis2** is a short projection directed posteriorly, while its **al** covers a broader range (as it is, the antero-ventral half of **gp9**). Viewed laterally, the **al** of **olis2** is slightly convex. This configuration possibly provides some degree of rotational freedom to **gs9** vs. **gp9** & **gp8** (interlocked by **olis1**, which extends more posteriorly than **olis2**, including its **rh**), using the **rh** of **olis2** as a slightly movable axis. This supposed ability would allow **gp8** postero-ventral teeth to be exposed (instead of concealed by **gs9**) and then used by the insect to appreciate the adequacy of substrate for oviposition. The **gp8** is only partially concealed by **gs9**.

###### 2.2.4 *Tettigonia viridissima* (Linnaeus, 1758) (schematized under ‘Tettigoniidae’ in Fig. 2G)

The observed configuration of the ovipositor valves conforms that described by Cappe de Baillon (*12*). Unlike assumed by Kluge (*14*; among others) we argue that the olistheter interlocking **gs9** and **gp8** (thereafter **olis3**) is not homologous with **olis2**. Firstly, a protrusion from **gs9** and directed towards **gp9** (viz., the characteristic features of **olis2**) occurs at various levels along the ovipositor. It is clearly distinct from another well-delimited olistheter (viz. **olis3**). Secondly, as stated by Kluge (*14*), the Anostostomatidae possibly represent an ‘intermediate’ stage is which a well-delimited **olis2** co-occurs with the premises of an **olis3**, in the shape of a projection of the ventral margin of **gs9** into **gp8**. If two olistheters occur (in addition to **olis1**), they cannot be homologous. It follows that there is an **olis3** besides **olis2**.

##### 2.3 Analysis of the mandibular mechanical advantage

Progression of mechanical advantage curves for the studied taxa are represented in fig. S9. Results of the PCA are summarized in tab. S3 and represented in fig. S10, including the phylogenetic PCA. Animated versions of the PCA represented in Fig. 3E are provided in the associated Dryad dataset (*58*).

#### 3 Insect species currently known to occur at Xiaheyan

Palaeoptera (24 sp.)

> Rostropalaeoptera (9 sp.)
>
> Palaeodictyoptera (4 sp.)
>
> *Namuroningxia elegans* Prokop & Ren, 2007
>
> *Sinodunbaria jarmilae* Li, Ren, Pecharová & Prokop, 2013
>
> *Xiaheyanella orta* Fu, Béthoux, Yand & Ren, 2015
>
> *Tytthospilaptera wangae* Liu, Béthoux, Yin & Ren, 2015
>
> Megasecopteromorpha (5 sp.)
>
> *Brodioptera sinensis* Pecharová, Ren & Prokop, 2015
>
> *Sinopalaeopteryx splendens* Pecharová, Prokop & Ren, 2015
>
> *Sinopalaeopteryx olivieri* Pecharová, Prokop & Ren, 2015
>
> *Namuroptera minuta* Pecharová, Prokop & Ren, 2015
>
> *Sinodiapha ramosa* Yang, Ren & Béthoux, 2020

Odonatoptera (6 sp.)

> *Shenzhousia qilianshanensis* Zhang & Hong, 2006
>
> *Oligotypus huangheensis* Ren, Nel & Prokop, 2008
>
> *Tupus orientalis* (Zhang, Hong & Su, 2012)
>
> *Erasipterella jini* (Zhang, Hong & Su, 2012)
>
> *Aseripterella sinensis* Li, Béthoux, Pang & Ren, 2013
>
> *Sylphalula laliquei* Li, Béthoux, Pang & Ren, 2013

Neoptera (17 sp.)

> Dictyoptera (5 sp.)
>
> *Qilianiblatta namurensis* Zhang, Schneider & Hong, 2013 *Kinklidoblatta youhei* Wei, Béthoux, Guo, Schneider & Ren, 2013 Undetermined sp.1
>
> Undetermined sp.2
>
> Undetermined sp.3

Grylloblattida (1 sp.)

> *Sinonamuropteris ningxiaensis* Peng, Hong & Zhang, 2005

Plecoptera (1 sp.)

> *Gulou carpenteri* Béthoux, Cui, Kondratieff, Stark & Ren, 2011

Archaeorthoptera (10 sp.)

> *Sinopteron huangheense* Prokop & Ren, 2007
>
> *Chenxiella liuae* Liu, Ren & Prokop, 2009
>
> *Longzhua loculata* Gu, Béthoux & Ren, 2011
>
> *Heterologus duyiwuer* Béthoux, Gu & Ren, 2012
>
> *Miamia maimai* Béthoux, Gu, Yue & Ren, 2012
>
> *Xixia huban* Gu, Béthoux & Ren, 2014
>
> *Protomiamia yangi* Du, Béthoux, Gu & Ren, 2017
>
> *Sinogerarus pectinatus* Gu, Béthoux & Ren, 2017
>
> *Phtanomiamia gui* Chen, Ren & Béthoux, 2020
>
> *Ctenoptilus frequens* sp. nov.

## SUPPLEMENTAL FIGURES

**Figure S1.**
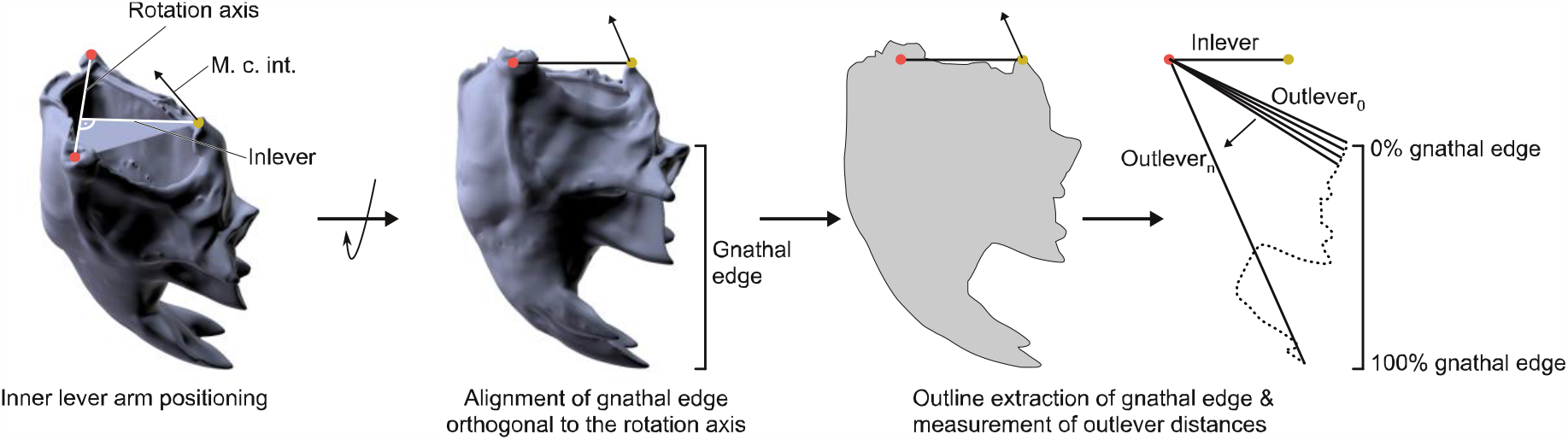
Workflow for the extraction of the mandibular mechanical advantage based on 3D models.

**Fig. S2.**
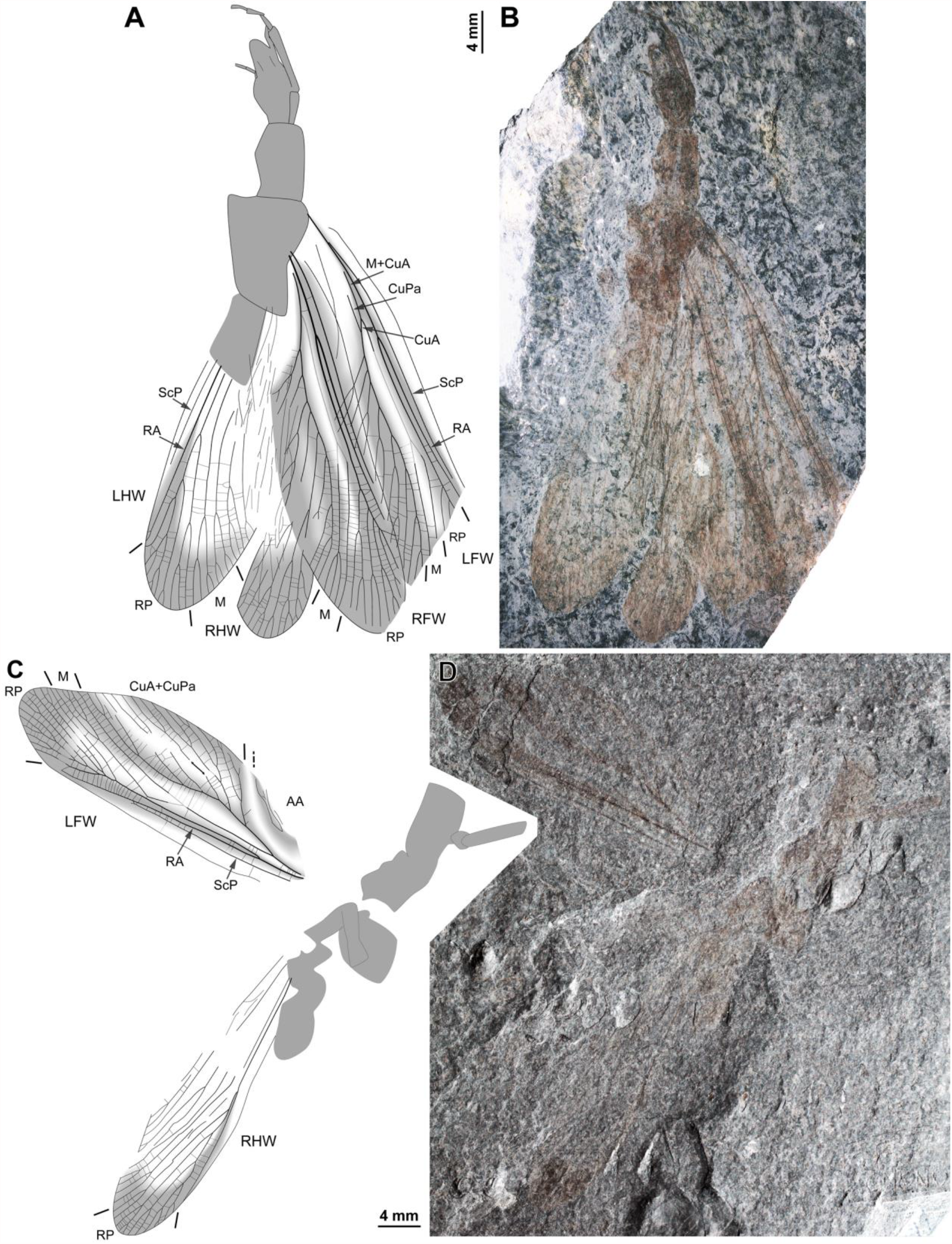
*Ctenoptilus frequens* sp. nov., specimens composed of fore- and hind wings in connection with body remains. (A–B) Specimen CNU-NX1-752; habitus, left forewing as positive imprint and right forewing and hind wings as negative imprints, (A) drawing and (B) photograph (composite). (C–D) Specimen CNU-NX1-738; habitus, right hind wing as positive imprints and left forewing as negative imprints, (C) drawing and (D) photograph (composite; slightly shifted vertically with respect to drawing).

**Fig. S3.**
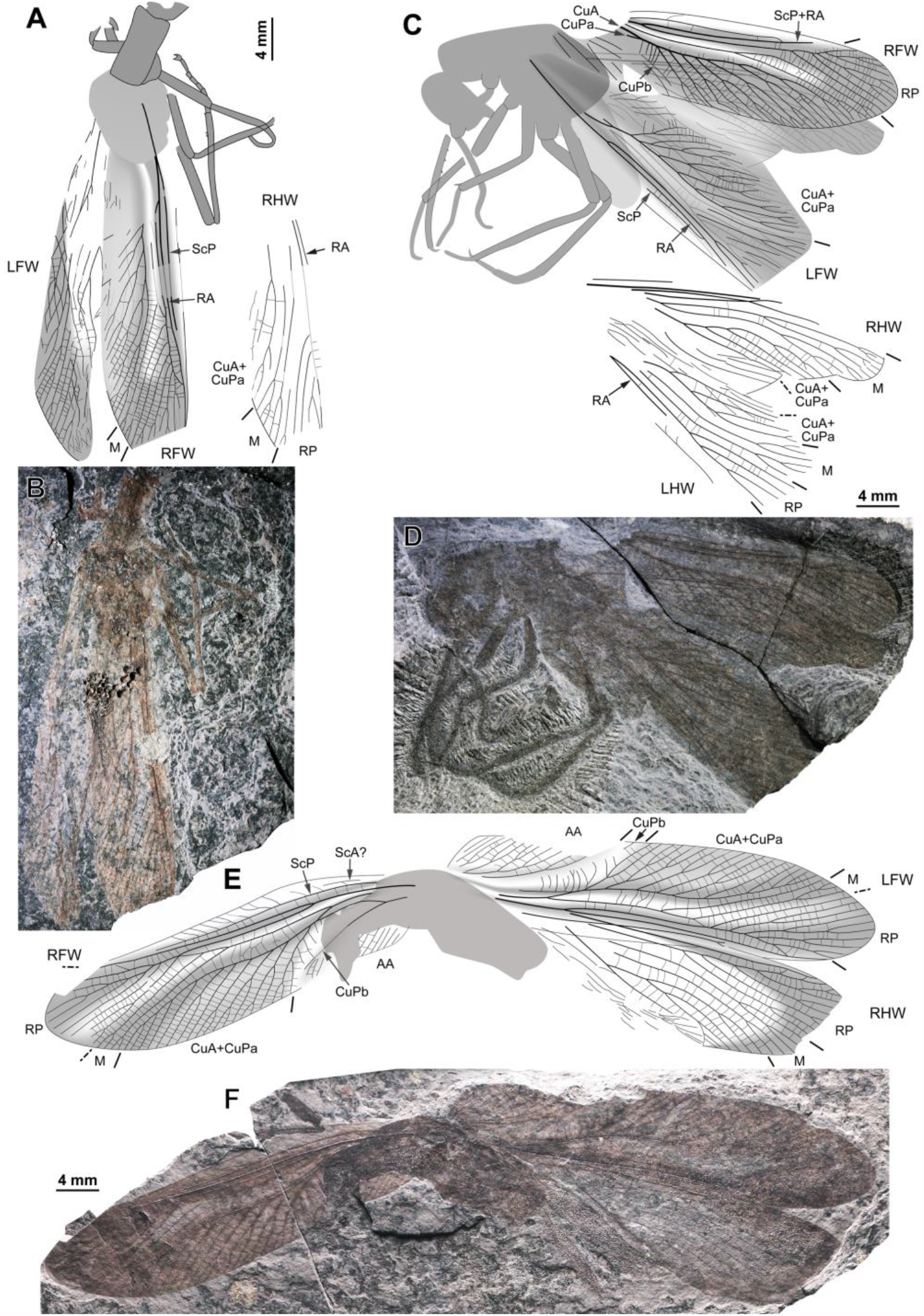
(preceding page). *Ctenoptilus frequens* sp. nov., specimens composed of fore- and hind wings in connection with body remains. (A–B) Specimen CNU-NX1-759; habitus, left hind wing as positive imprint and right wings as negative imprints, (A) drawing (for clarity, drawing of right hind wing venation duplicated and relocated, original location in light grey on complete drawing) and (B) photograph (composite). (C–D) Specimen CNU-NX1-750; habitus, all wings as negative imprints, (C) drawing (for clarity, drawing of hind wings venation duplicated and relocated, original location in light grey on complete drawing) and (D) photograph (composite). (E–F) Specimen CNU-NX1-731; habitus, left forewing as positive imprint and right forewing and right hind wing as negative imprints, (E) drawing and (F) photograph (composite).

**Fig. S4.**
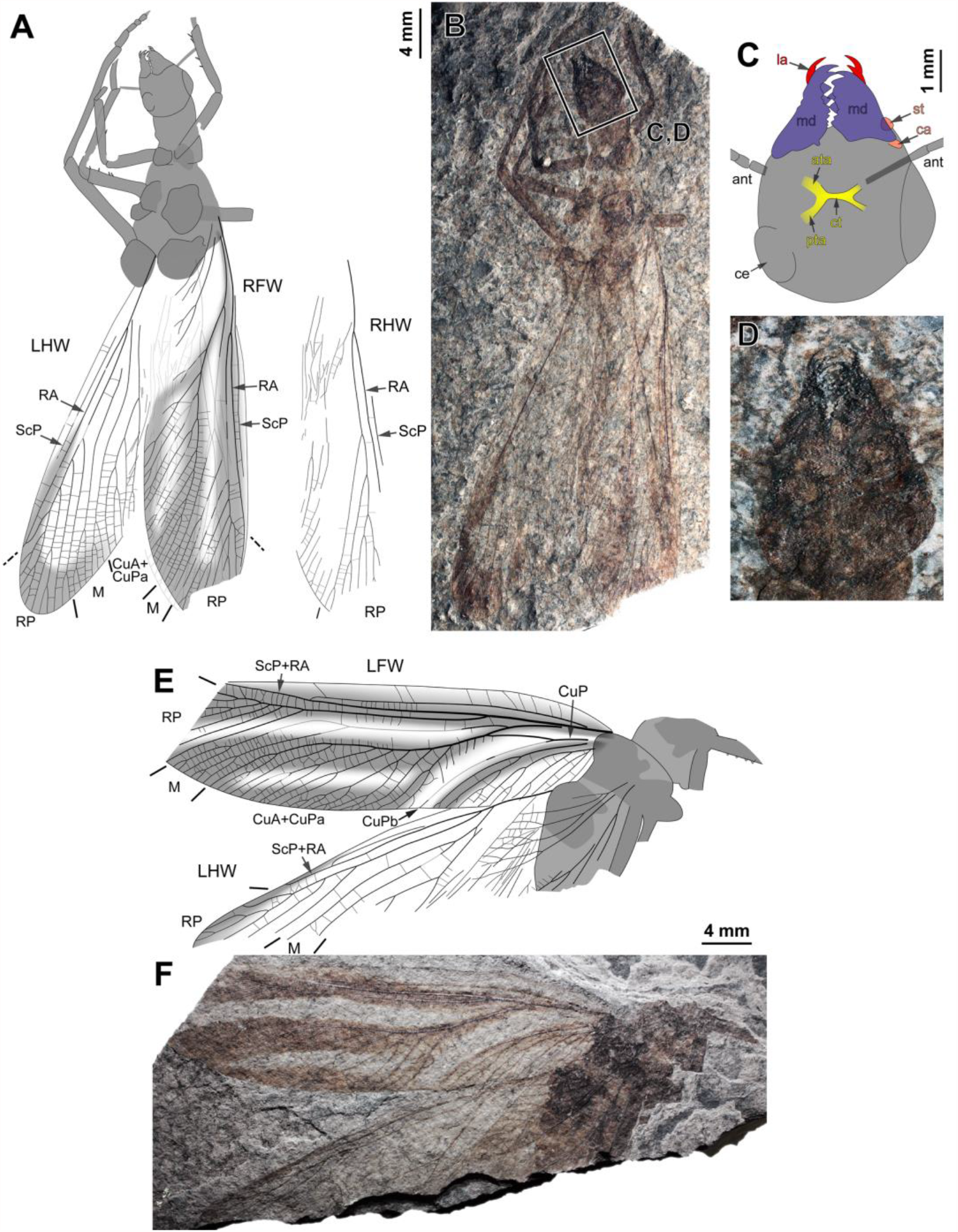
*Ctenoptilus frequens* sp. nov., specimens composed of fore- and hind wings in connection with body remains. (A–D) Specimen CNU-NX1-747; (A–B) habitus, all wings as negative imprints, (A) drawing (for clarity, drawing of right hind wing venation duplicated and relocated, original location in light grey on complete drawing) and (B) photograph (composite); and (C–D) details of head (location as indicated in B), polarity unclear, (C) color-coded interpretative drawing and (D) photograph (composite). Color-coding: red, lacina (**la**); salmon, cardinal and stipital sclerites (**ca** and **st**, respectively); dark blue-purple, mandible (**md**); yellow, tentorium, including anterior tentorial arm (**ata**), posterior tentorial arm (**pta**) and corpotentorium (**ct**). Other indications: **ant**, antenna; **ce**, composite eye. (E–F) Specimen CNU-NX1-741; habitus, all wings as positive imprints, (E) drawing and (F) photograph (composite).

**Fig. S5.**
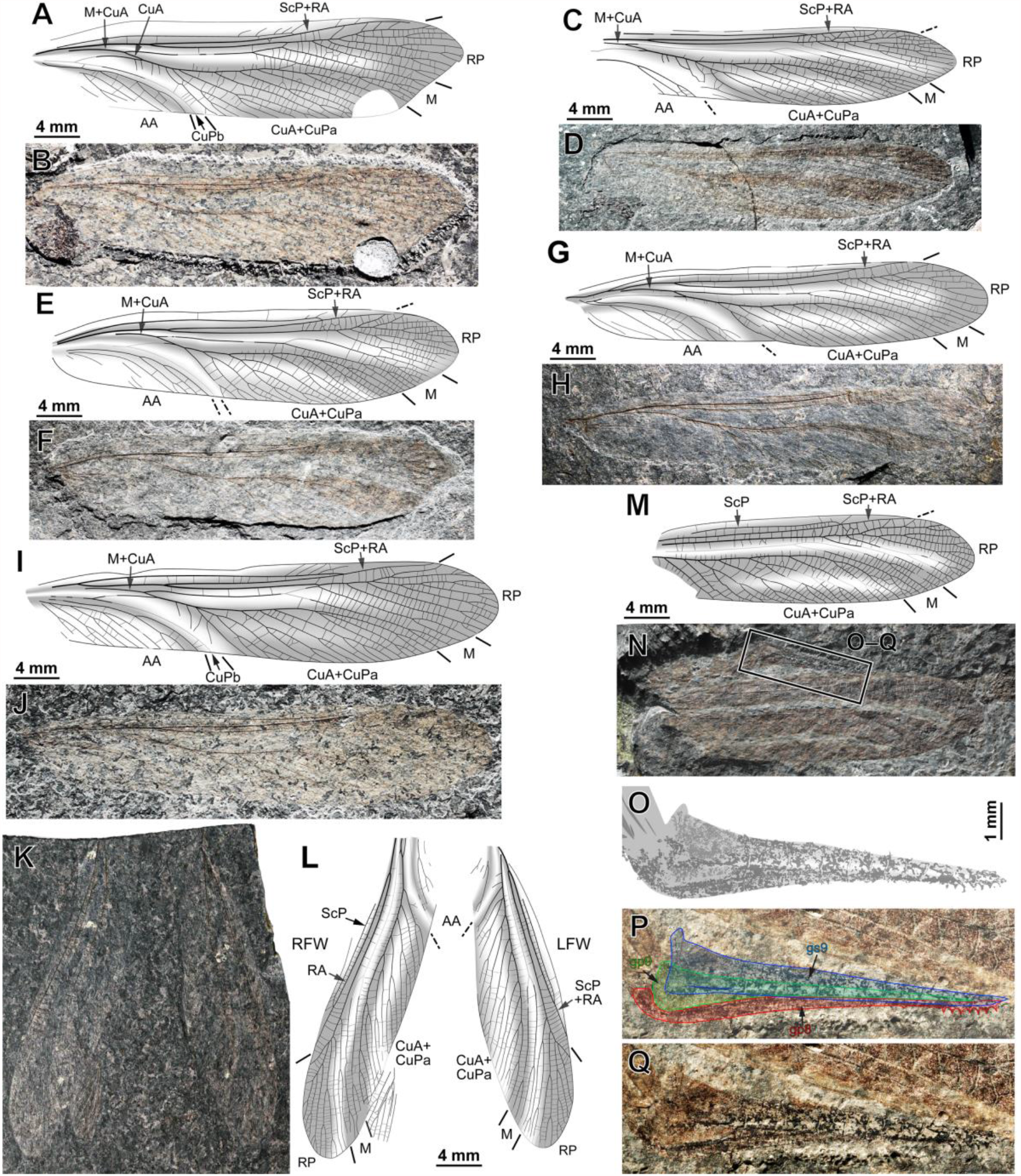
(preceding page). *Ctenoptilus frequens* sp. nov., specimens composed of forewings, isolated or by pair, and forewing and ovipositor. (A–B) Specimen CNU-NX1-748; right forewing, negative imprint, (A) drawing and (B) photograph (composite, flipped horizontally, light-mirrored). (C–D) Specimen CNU-NX1-732; right forewing, positive imprint, (C) drawing and (D) photograph (composite). (E–F) Specimen CNU-NX1-757; right forewing, negative imprint, (E) drawing and (F) photograph (composite, flipped horizontally, light-mirrored). (G–H) Specimen CNU-NX1-758; left forewing, negative imprint, (G) drawing and (H) photograph (composite). (I–J) Specimen CNU-NX1-744; right forewing, negative imprint, (I) drawing and (J) photograph (composite, flipped horizontally, light-mirrored). (K–L) Specimen CNU-NX1-751; forewing pair, both as negative imprints, and apical fragment of a hind wing, (K) drawing and (L) photograph (composite). (M– Q) Specimen CNU-NX1-743; (M–N) habitus, right forewing, positive imprint, (M) drawing and (N) photograph (composite); and (O–Q) details of ovipositor (location as indicated in N), polarity unknown, (O) drawing and (P– Q) photographs, (P) with color-coded interpretative drawing and (Q) without (composite, flipped horizontally).

**Fig. S6.**
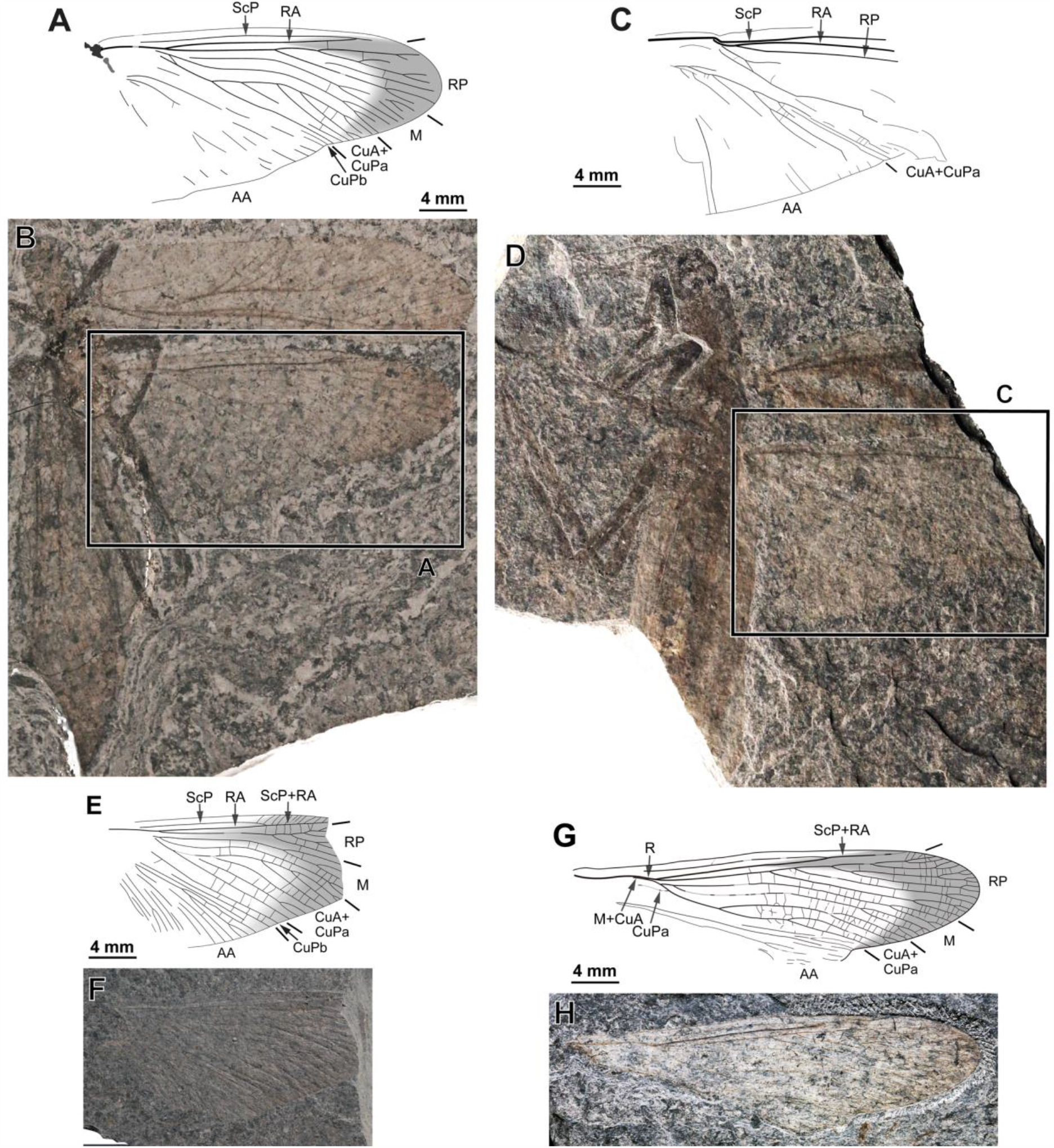
*Ctenoptilus frequens* sp. nov., specimens composed of well-exposed hind wings in connection with body remains or isolated. (A–B) Specimen CNU-NX1-198; (A) drawing of right hind wing (location as indicated in B) and (B) photograph of habitus (composite, flipped horizontally), left forewing as positive imprints and left hind wing and right wings as negative imprints. (C–D) Specimen CNU-NX1-740; (C) drawing of right hind wing; (D) photograph for habitus (composite, flipped horizontally, light-mirrored), right wings as positive imprints. (E–F) Specimen CNU-NX1-199; right hind wing, positive imprint, (E) drawing and (F) photograph (composite). (G–H) Specimen CNU-NX1-753; left hind wing, negative imprint, (G) drawing and (H) photograph (composite).

**Fig. S7.**
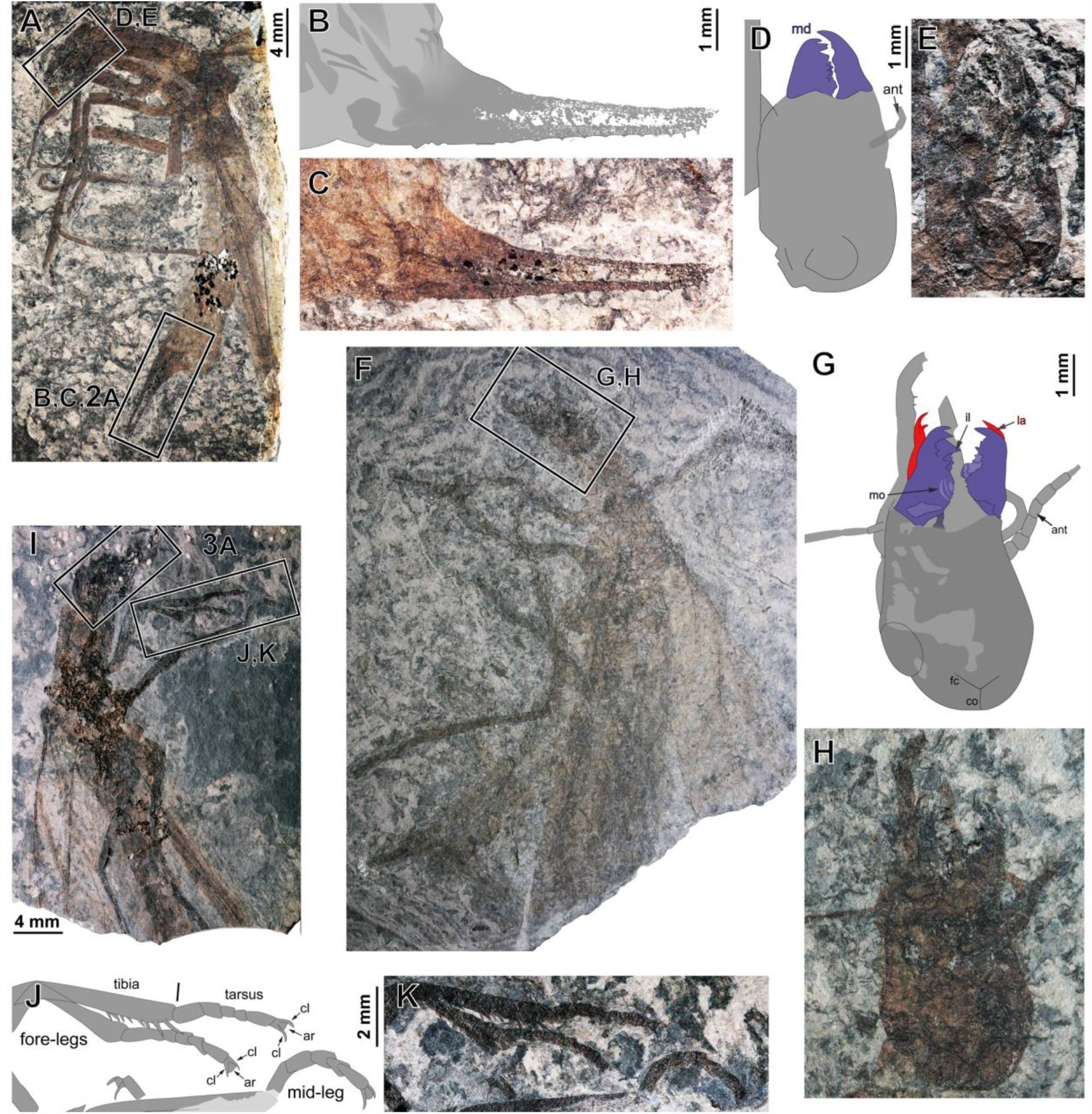
*Ctenoptilus frequens* sp. nov., specimens composed of body remains including well-preserved head, legs and/or ovipositor. (A–E) Specimen CNU-NX1-749; (A) photograph of habitus (composite), left forewing as positive imprint; (B–C) details of ovipositor (location as indicated in A; to be compared with main document Fig. 2C), polarity unclear, (B) drawing and (C) photograph (composite); and (D–E) details of head (location as indicated in A), (D) color-coded interpretative drawing and (E) photograph (composite). (F–H) Specimen CNU-NX1-756; (F) photograph of habitus (composite); and (F–H) details of head (location as indicated in F), imprint polarity unclear, (G) color-coded interpretative drawing and (F) photograph (composite). (I–K) Specimen CNU-NX1-754; (I) photograph of habitus (composite; frame delimiting head indicating the location of main document Fig. 3A–B), (J–K) details of distal portions of fore-legs and a mid-leg (location as indicated in I), (J) drawing and (K) photograph (composite).

**Fig. S8.**
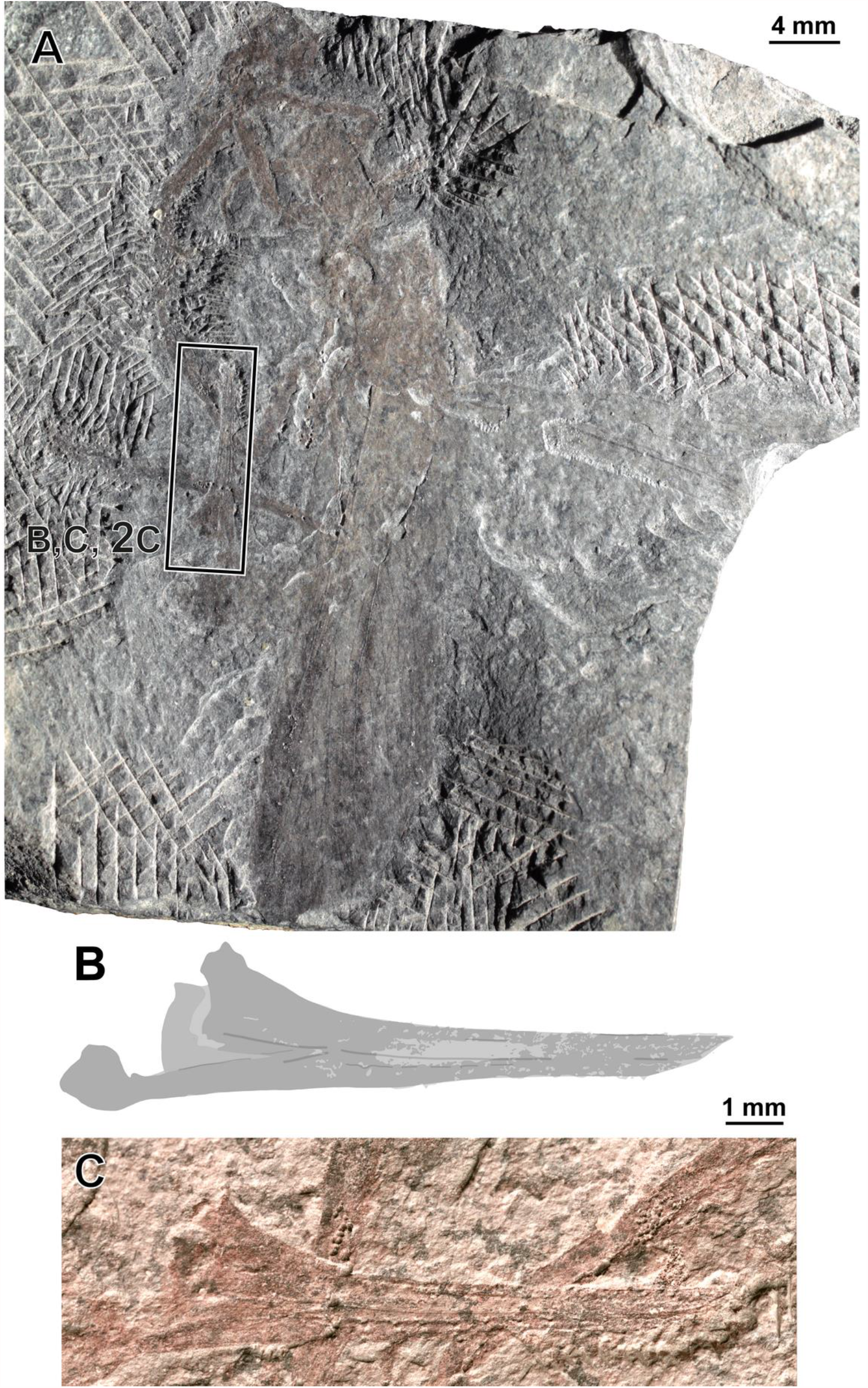
*Ctenoptilus frequens* sp. nov., specimen CNU-NX1-742. (A) Photograph of habitus (composite), right forewing as positive imprint, flipped horizontally, and (B–C) details of ovipositor (location indicated in A; to be compared with main document Fig. 2E), (B) drawing and (C) photograph (light-mirrored).

**Fig. S9.**
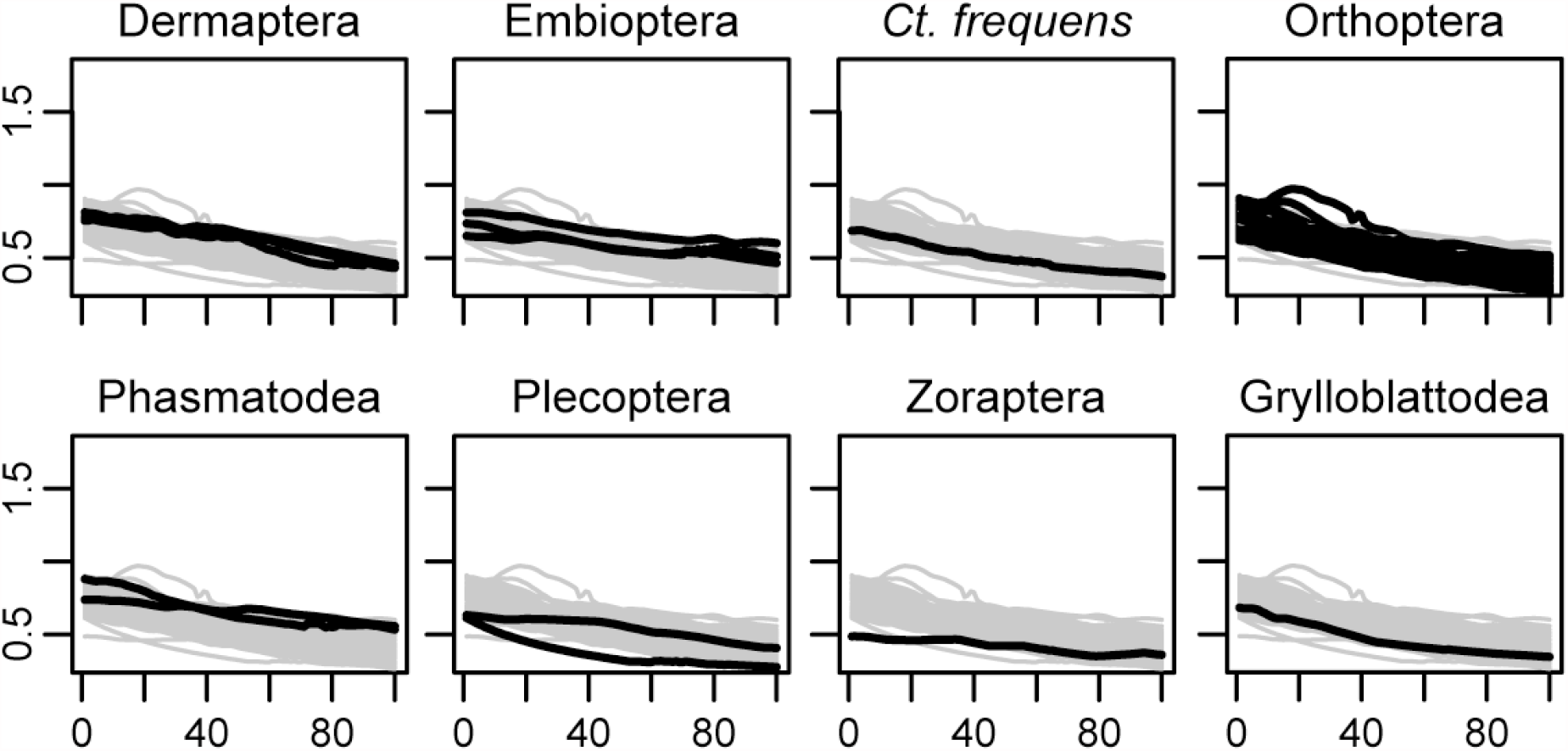
Progression of mechanical advantage curves for the studied taxa. x-axis = % tooth row; y-axis=MA.

**Fig. S10.**
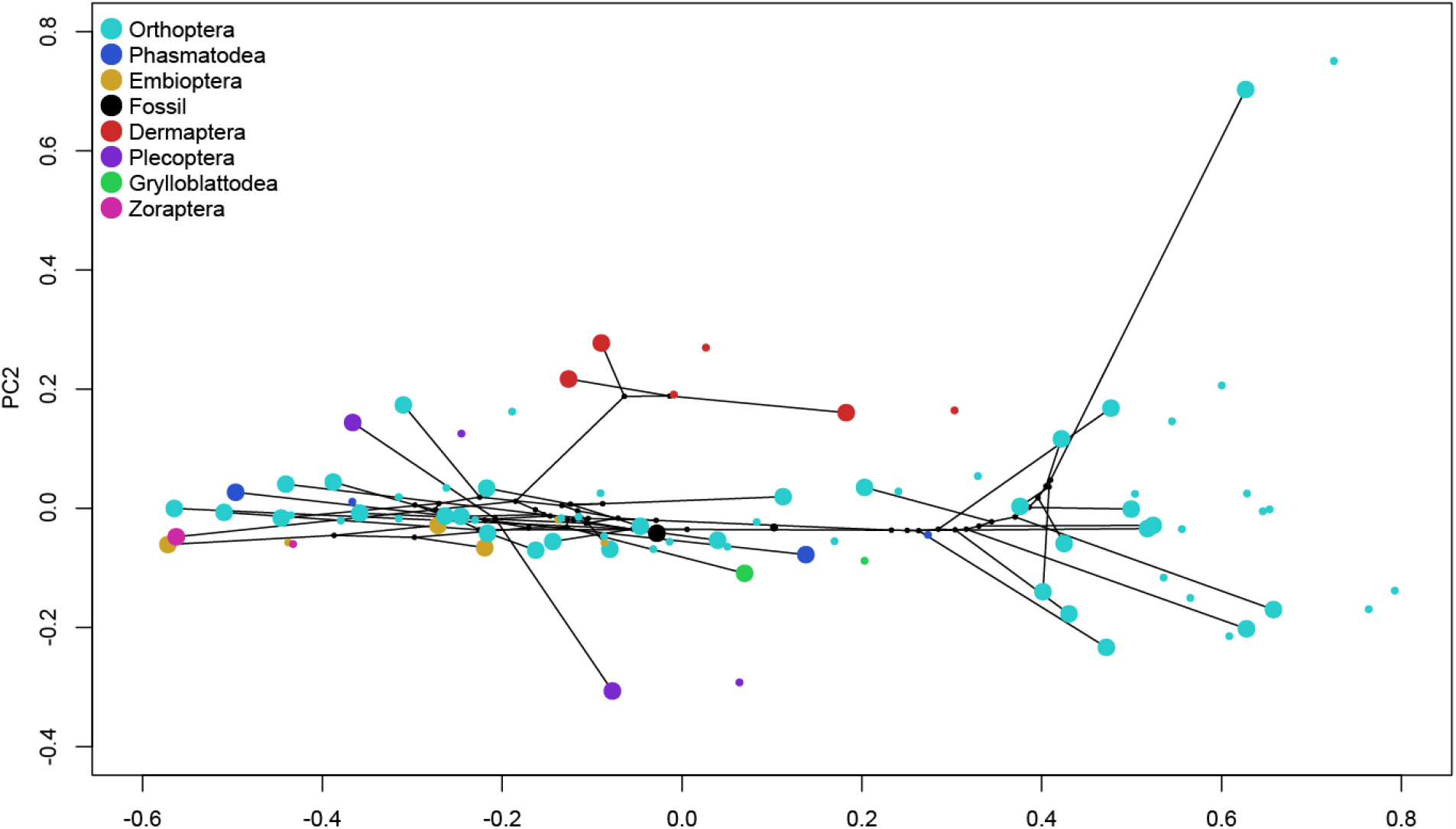
Results of the principal component analysis of the mandibular mechanical advantage for the first two PCs together with results for the first two PCs after phylogenetic signal correction. Large dots, distribution of species in PC space uncorrected for phylogenetic signal; small dots, distribution of species in PC space corrected for phylogenetic signal. Although phylogenetic signal was significant, differences do not affect the relative position of the sampled species to each other in PC space.

## SUPPLEMENTAL TABLES

**Table S1.**
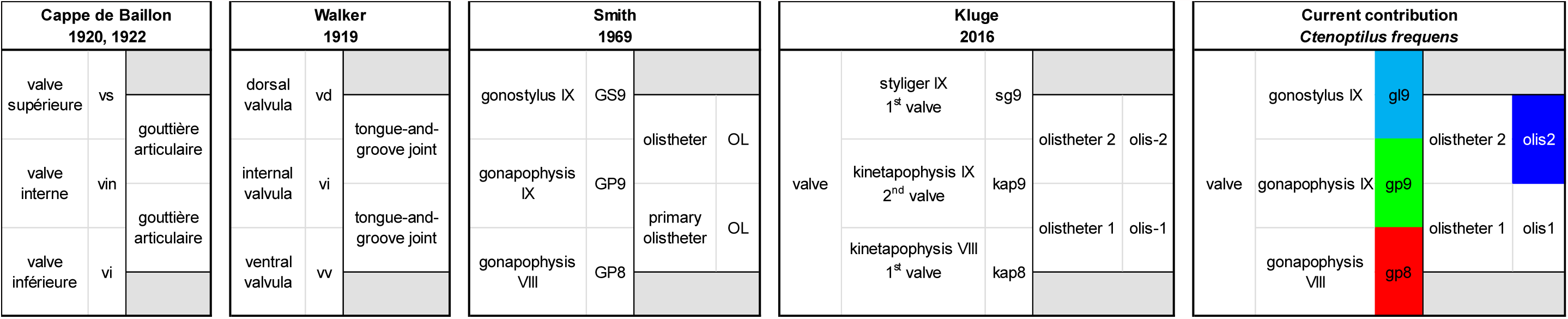
Correspondence between terminologies applied to polyneopteran insect ovipositors.

**Table S2.**
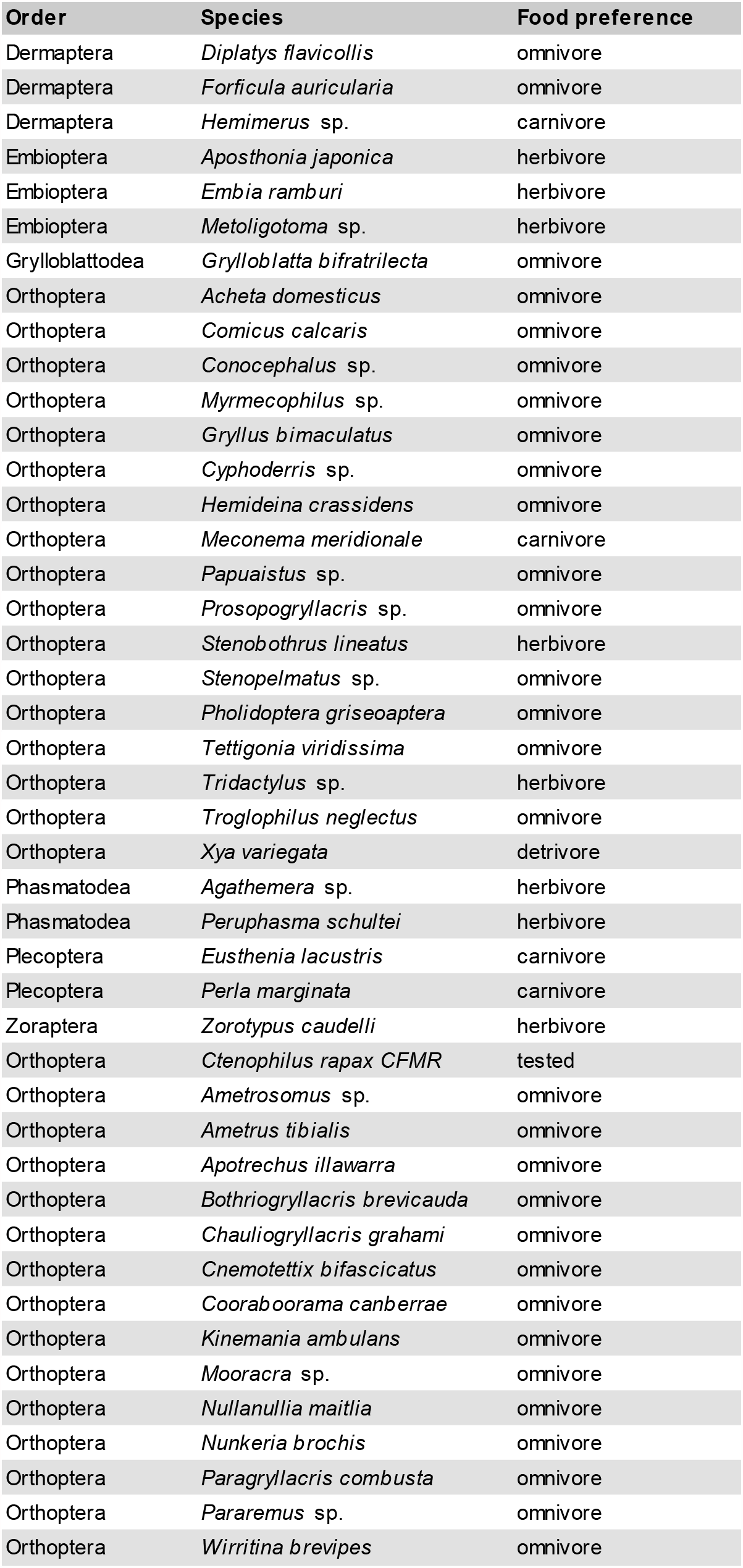
Food preference of polyneopteran species included in the MA Principal Component Analyses.

**Table S3.**
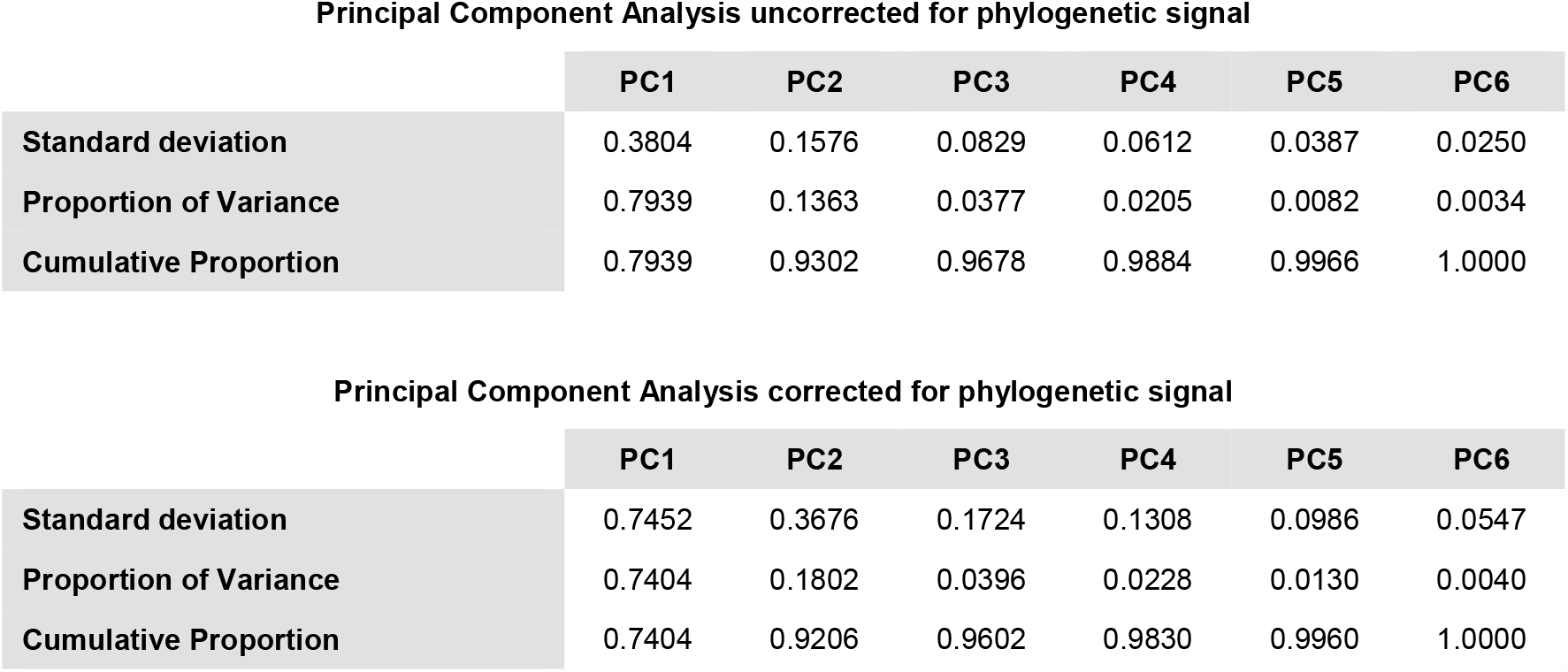
Importance of components of Principal Component Analyses of MA.

**Principal Component Analysis uncorrected for phylogenetic signal**
|  | **PC1** | **PC2** | **PC3** | **PC4** | **PC5** | **PC6** |
| --- | --- | --- | --- | --- | --- | --- |
| **Standard deviation** | 0.3804 | 0.1576 | 0.0829 | 0.0612 | 0.0387 | 0.0250 |
| **Proportion of Variance** | 0.7939 | 0.1363 | 0.0377 | 0.0205 | 0.0082 | 0.0034 |
| **Cumulative Proportion** | 0.7939 | 0.9302 | 0.9678 | 0.9884 | 0.9966 | 1.0000 |

|  | **PC1** | **PC2** | **PC3** | **PC4** | **PC5** | **PC6** |
| --- | --- | --- | --- | --- | --- | --- |
| **Standard deviation** | 0.7452 | 0.3676 | 0.1724 | 0.1308 | 0.0986 | 0.0547 |
| **Proportion of Variance** | 0.7404 | 0.1802 | 0.0396 | 0.0228 | 0.0130 | 0.0040 |
| **Cumulative Proportion** | 0.7404 | 0.9206 | 0.9602 | 0.9830 | 0.9960 | 1.0000 |

